# Actinium-225 Pretargeted Radioimmunotherapy against Colorectal Cancer using Self-Assembling and Disassembling Antibodies

**DOI:** 10.64898/2026.11.18.767407

**Authors:** Nicole Aguirre, Navya Vemula, Sang-Gyu Lee, Sebastian E. Carrasco, Renata Mammone, Hong-fen Guo, Sebastian Chung, Edward K. Fung, Alex Le Roux, Jorge A. Carrasquillo, Karina Leung, Michelle Kim, Hong Xu, Yi Feng, Irene Cheung, Andrea Cercek, Michael B. Foote, Georgios Karagkounis, Steven M. Larson, Sarah M. Cheal, Darren Veach, Garrett M. Nash, Nai-Kong V. Cheung

**Author notes:** Corresponding Author: Nai-Kong Cheung MD PhD.

## Abstract

Colorectal cancer (CRC) remains a leading cause of cancer deaths worldwide. GPA33, expressed on 95% of CRC, has been extensively targeted by IgG-attached payloads in conventional radioimmunotherapy (RIT), but toxicity was dose-limiting. A 2-step pretargeted radioimmunotherapy (PRIT) regimen using Self-Assembling-Disassembling antibody (SADA) specific for GPA33 (10 mg/kg) intravenous (IV) in step #1 and 24 h later [^86^Y]Y-ABD (9 nmole) IV in step #2, avoided normal gut uptake in non-human primates. Dosimetry results based on tissue counting in murine models using GPA33- SADA IV in step #1 and [^225^Ac]Ac-Pr in step #2, were 30.88, 0.1 and 2.56 Gy/μCi (Gy/37 kBq; RBE= 1), respectively for subcutaneous (sq) SW1222 tumor, blood and kidney, demonstrating favorable therapeutic ratios (309 for blood; 12 for kidney). Established sq SW1222 xenografts treated with ^225^Ac-SADA-PRIT showed dose dependent improvement in median survival (MS). Using one cycle of 2 doses 7 days apart (74kBq + 74kBq of ^225^Ac), majority of mice had complete tumor volumetric responses of sq SW1222, LS174T, LOVO and intraperitoneal SW1222 xenografts. Complete response was confirmed histologically at 180 days, accompanied by statistically significant improvements in MS. All treatments were well tolerated, with transient reversible weight loss (<15%), minimal hematologic, and no acute (within 30 days post ^225^Ac) chemical toxicities. At necropsy, kidneys showed minimal to marked subclinical tubular injury.

The ability of a single cycle of ^225^Ac-SADA-PRIT to induce major CRC response and durable remission with minimal acute toxicities in multiple xenograft models while avoiding on-target off-tumor gut toxicity is encouraging, although subclinical histologic renal and ovarian damage remain a concern.

## INTRODUCTION

Colorectal cancer (CRC) is the third most commonly diagnosed cancer and the second leading cause of cancer-related mortality worldwide.^1,2^ In 2020, more than 1.9 million new CRC cases were diagnosed, and nearly 0.9 million deaths were attributed to the disease. Nearly 20–30% present with metastatic disease, where treatment options remain limited and 5-year survival is <20%. ^3^

Glycoprotein A33 antigen (GPA33) is a transmembrane glycoprotein highly expressed in over 95% of human CRCs and expression limited to the normal gastrointestinal epithelium. As a tumor target, GPA33 exhibits minimal internalization or shedding, thus prolonging tumor surface retention of bound antibodies. Normal tissue expression can be overcome by physiologic gut epithelium turnover to reduce drug exposure.^4^ Radioimmunotherapy (RIT) delivers “liquid radiation” via anti-tumor targeted antibodies conjugated with radioisotopes, making it particularly well-suited for treating metastatic disease.^5^ Despite the favorable attributes of GPA33, IgG based drug delivery systems achieved poor therapeutic indices (TI) and a narrow therapeutic window (TW), leading to unintended bystander toxicity in critical organs (e.g. marrow, kidney, liver), as well as on-target, off-tumor (e.g. gut) side effects. GPA33 seems like an ideal target for 2-step pretargeted radioimmunotherapy (PRIT), which requires sufficient antibody retention on the tumor surface during step #1, and rapid normal tissue clearance prior to administration of the radioactive payload in step #2.

We recently described a novel self-assembling and disassembling (SADA) bispecific antibody (BsAb) platform for 2-step PRIT to safely deliver tumoricidal doses of alpha (α)- or beta (β)-emitting radiation (Figure 1). The SADA platform is built with basic tandem single-chain variable fragments (scFvs): one scFv targeting the tumor antigen and the other scFv recognizing the BnDOTA(La) (La = Lanthanide) metal complex. A p53-derived tetramerization domain enables self-assembly into stable tetramers (220 kDa) to extend half-life from minutes to hours, while allowing dissociation of unbound antibodies into monomers (55 kDa) for rapid renal clearance. Compared to IgG, the fast clearance of SADA limits its transit into luminal epithelial cells while reducing its immunogenicity.

**Figure 1:**
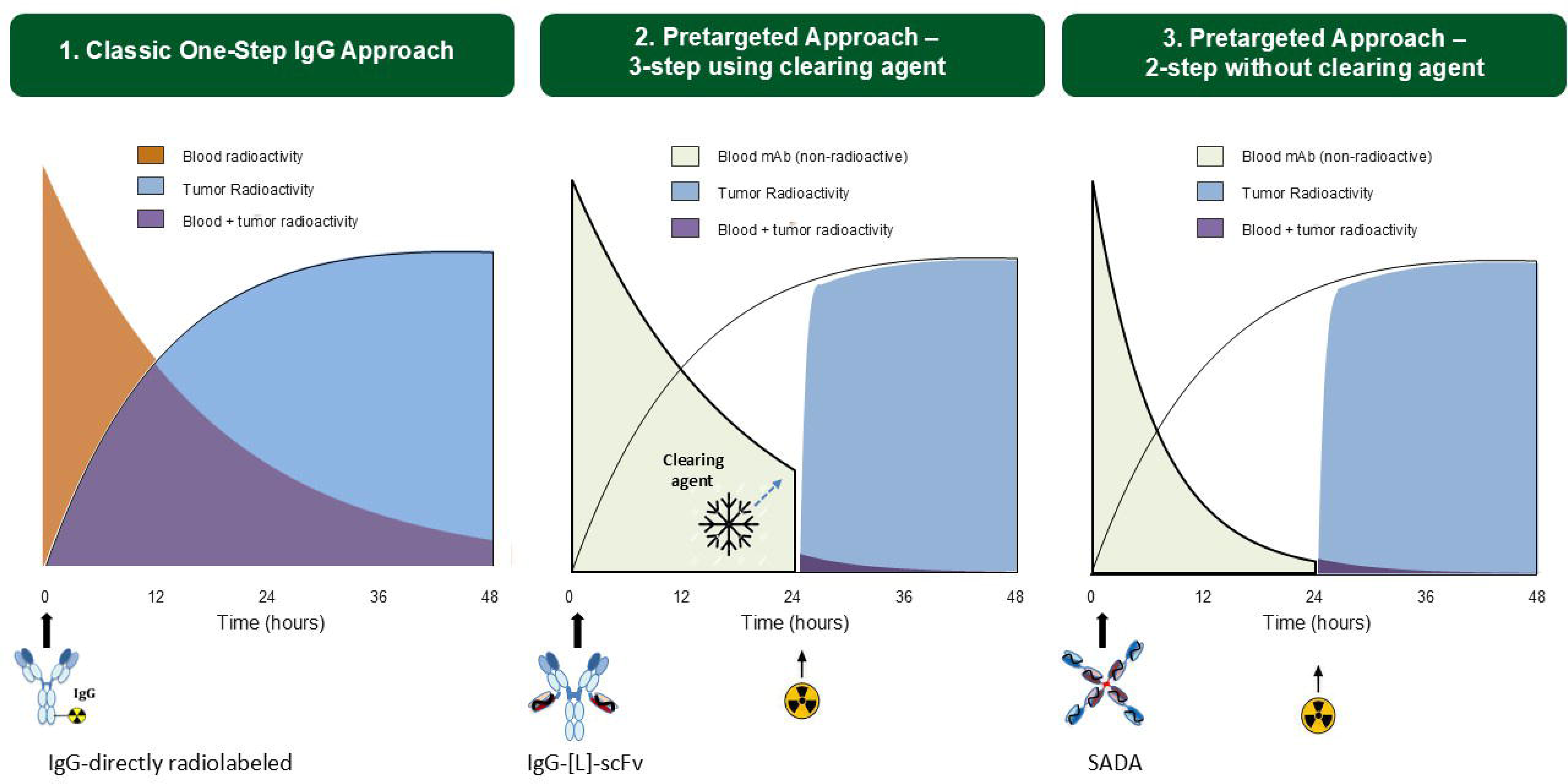
Past to future of radioimmunotherapy: **(A)** Conventional radioimmunotherapy using IgG in a single step fashion; **(B)** DOTA-PRIT with IgG-scFv in 3-step PRIT; **(C.)** SADA in 2-step PRIT. SADA bispecific antibodies (BsAbs) have been developed successfully against a broad spectrum of human cancer targets, including GD2, GPA33, and CD38.

^225^Ac is a long-lived (t_1/2_ = 9.92 d) α-emitter. Each α particle is composed of two protons and two neutrons, carries 5.8 to 8.4 MeV energy with a charge of +2, and penetrates 50 to 100 µm in tissues. Its high linear energy transfer creates difficult-to-repair double- strand DNA breaks and is relatively independent of oxygen tension, distinguishing it from conventional radiation modalities where tumor hypoxia has been a major cause of treatment failure. The potency and precision of α particles have been recognized for over a century, though harnessing them safely for medical use has proven challenging.^6,7^ More recently, pivotal clinical trials using ^225^Ac in radiopharmaceutical therapy (RPT) have shown encouraging results.^8^ These advances come as supply chain challenges for alpha emitting radioisotopes are being addressed. Using the novel SADA antibody platform against GPA33 in CRC, we tested the potential of alpha particle nanogenerator ^225^Ac in 2-step alpha PRIT to overcome the two main hurdles of insufficient TI/TW and on-target off-tumor limitations to establish a curative and safe approach to CRC in preclinical models.

## MATERIALS AND METHODS

### PRIT Reagents

SADA-BsAb proteins were prepared as described previously.^9^ Briefly, expression was performed using the Expi293 Expression System (Invitrogen) and purified by affinity chromatography (GE Healthcare, Ni-NTA Gravitrap, cat# 11003399). Biochemical purity of candidate SADA were determined by size exclusion chromatography-HPLC (SEC- HPLC).

The ^225^Ac used in this research was supplied by the United States Department of Energy Office of Science by the Isotope Program in the Office of Nuclear Physics (source: ^229^Th stockpile). Proteus (molecular weight: 1,350 g/mol), the chelator for ^225^Ac, consists of an empty DO3A separated by a PEG4 linker to a ^175^Lu-complex of 2-benzyl- DOTA. ^225^Ac-Proteus was prepared using previously described methods to a specific activity of 1.05 Ci/g (5.2 x 10^4^ GBq/mol). ^225^Ac-activity measurements were performed at secular equilibrium using a CRC-15R radioisotope calibrator (Capintec) set at 775. The displayed activity value was multiplied by a factor of 5, and samples were positioned at the bottom center of the well for consistent measurement.

### Animal Studies

All animal experiments were performed in accordance with protocols approved by the Institutional Animal Care and Use Committee of Memorial Sloan Kettering Cancer Center (MSKCC; New York, NY) following NIH guidelines for animal welfare. Animal experiments used female athymic nude mice (strain: Hsd:Athymic Nude-Foxn1nu, Envigo; aged 6–8 weeks; average weight for 6-week-old and 8-week-old animals: 18.6 and 21.0g, respectively). For all injections, mice were gently warmed with a heat lamp and placed on a restrainer. Their tails were sterilized with alcohol pads and single-bolus injections were placed into the lateral tail vein. All mice from the same treatment groups were cohoused in the same cage. Blinding of treatment or experimental measurements was not performed.

In vivo studies utilized SW1222 (Ludwig Institute for Cancer Immunotherapy; New York, NY), LOVO, or LS174T CRC cell lines. Mice were subcutaneously inoculated with 5 x 10^6^ cells suspended in 100 µL of a 1:1 mixture of culture media and Matrigel (Corning), using a sterile syringe fitted with a 28-gauge needle. Unless otherwise noted, experiments were initiated 7-12 d post-inoculation, when tumor volumes (calculated assuming ellipsoidal geometry) reached approximately 300–500 mm^3^.

### Imaging and Dosimetry in Nonhuman Primates

Cynomolgus monkeys were used to evaluate potential on-target, off-tumor gastrointestinal uptake of GPA33-targeted PRIT. Animals received intravenous (IV) GPA33 IgG-scFv or GPA33-SADA bispecific antibodies (0.2 mg/kg), followed 24 hours later by IV [^86^Y]Y-aminobenzylDOTA ([^86^Y]Y-ABD; 9 nmol). Serial PET/CT imaging and tissue uptake analyses were performed to compare gastrointestinal biodistribution of IgG- and SADA-based constructs. An IgG-scFv variant with silenced FcRn affinity (H310A) was also evaluated. Monkey anti-human antibody responses were measured longitudinally by in-house ELISA over 4 months to assess immunogenicity.

For longitudinal imaging, two healthy male Mauritian cynomolgus macaques (Macaca fascicularis; PrimGen, Inc.), aged 4-5 years and weighing ∼4 kg, were studied at the Citigroup Biomedical Imaging Center, Weill Cornell Medicine, using a Biograph64 mCT PET/CT scanner (Siemens Medical Solutions). Animals were sedated and administered GPA33-SADA (0.2 mg/kg or 10 mg/kg IV) or saline sham 24 hours before imaging. For PET/CT, animals were maintained under isoflurane anesthesia, and [86Y]DOTA or [86Y]BnDOTA (37–74 MBq; 1–2 mCi) was administered IV via a saphenous PICC, followed by a saline flush. PET acquisition began at radiotracer administration, with a repeat 30-minute PET/CT scan performed 24-hours later without additional radiotracer. PET data were reconstructed using ordered-subset expectation maximization with attenuation, scatter, and Y-86 prompt-gamma corrections. Volumes of interest (VOIs) were drawn on co-registered PET/CT images for organs of interest using Hermes Hybrid Viewer (Hermes Medical Solutions), and decay-corrected activity within each VOI was expressed as percentage injected dose (%ID).

All studies using cynomolgus macaques were approved by the IACUC of MSKCC. Animals were maintained in accordance with the USDA Animal Welfare Act and Regulations and the Guide for the Care and Use of Laboratory Animals. The animal care and use program at MSK is USDA-registered, maintains Public Health Services Assurances, and is fully accredited by AAALAC International.

### Tissue Biodistribution Analysis and Dosimetry

For biodistribution assay following radiohapten injection, mice were euthanized by CO_2_ (g) asphyxiation. Tumor and selected organs were harvested, rinsed with PBS, air- dried, weighed, and radioassayed by gamma scintillation counting (Perkin Elmer Wallac Wizard 3”). Count rates were background- and decay-corrected, converted to activity using a system calibration factor specific for the isotope, normalized to the administered activity, and expressed as percentage injected dose per gram of tissue (%ID/g).

For ^225^Ac, each sample was counted for up to 10 minutes, 24 h after collection, once secular equilibrium was reached, using a 150 to 600 keV energy window.

### Therapy Studies

Female nude mice were implanted with tumors at the right flank 9-12 days before treatment. Tumor-bearing mice were randomized by tumor volume before treatment. Established tumors were treated with GPA33-SADA (TC170) on day -3 and with [^225^Ac]Ac-Pr (74 kBq) on day 0. One cycle consisted of two doses given 7 days apart, to assure full clearance of antibody and chelator prior to a second dose. Mice treated with three-step IgG-scFv (BC105 (n = 10, 23.6 ± 32.1 mm^3^); BC177 (n = 8, 26.1 ± 29.3 mm^3^)) were treated on day 0 and received 25 µg of a DOTA-dendrimer clearing agent 4 hours prior to administration of [^225^Ac]Ac-Pr.^10^ Controls included antibody only or administration of radioisotope [^225^Ac]Ac-Pr (74 kBq) only. No significant difference was observed among starting tumor volume (one-way ANOVA; P >0.5 for all experiments).

Treated mice were monitored daily and weighed up to twice weekly for evidence of treatment-induced toxicity. Tumors were measured manually up to twice weekly using digital calipers and tumor volumes were calculated assuming ellipsoidal geometry.

Animals were monitored for toxicity (weight loss of 25%, petechiae, tumor ulceration or tumor size >2000 mm^3^). At ∼180 days post treatment, all survivors were evaluated (with age-matched littermates for reference) for long-term radiation-induced histologic organ damage.

### Anatomic and Clinical Pathology for Toxicology Assessment

To assess acute hematologic toxicity during therapy, blood was collected via retroorbital blood draws in tubes containing EDTA at baseline, just prior to treatment start, as well as d14, d21 or d28, and periodically thereafter (n = 3 mice/point). Hematology and serum biochemistry were performed at the Laboratory of Comparative Pathology (LCP; a shared resource of MSKCC, the Rockefeller University, and Weill Cornell Medicine).

Complete blood counts with differentials were obtained using an automated hematology analyzer (IDEXX Procyte DX, Columbia, MO). Complete blood count values were compared to LCP’s in-house reference values from experimentally naïve athymic nude mice. Serum chemistry was performed by the LCP on a Beckman Coulter AU680 analyzer (Beckman Coulter Life Sciences, Brea, CA).

Mice submitted for postmortem examination were euthanized by CO2 asphyxiation and cardiac exsanguination. Complete or partial necropsies were performed at the Laboratory of Comparative Pathology (MSK, the Rockefeller University, and Weill Cornell Medicine). Representative sections from the tumors and major organs were fixed in 10% neutral-buffered formalin, processed in alcohol and xylene, embedded in paraffin, sectioned (5-μm-thick) and stained with hematoxylin and eosin. The skull, spinal column, sternum, and hindlimb were decalcified in a formic acid and formaldehyde solution (Surgipath Decalcifier I, Leica Biosystems, Wetzlar, Germany) before processing. Tissues were examined by board-certified veterinary pathologists (SEC, RM). Representative histological images of tissues were captured using a brightfield BX45 microscope equipped with a DP26 camera and cellSens (version 1.18) Dimension software (Olympus America, Center Valley, PA).

### Data Reporting and Statistical Analysis

Statistical analysis was performed using GraphPad Prism software. One-way ANOVA with Dunnett’s multiple comparison tests was used to compare areas under the curve for tumor growth experiments. Taking into account the effect size that represents the maximum difference in any two groups, n = 5 per group minimum was needed to detect a difference with 80% power and 5% level of significance.

## RESULTS

### Blood Clearance of GPA33-SADA and [^225^Ac]Ac-Pr

We first investigated the pharmacokinetics (PK) of the individual pretargeting components (GPA33 SADA and 225Ac-proteus).^11^ 250 μg (1 nmole, approximately 10 mg/kg GPA33-SADA dose) gave peak plasma concentrations (Cmax) above 200 μg/mL within 30 minutes of injection. A trough level of <1μg/mL was achieved at 48-80 hours

post injection in nude mice. Below this plasma concentration, 2 μCi (74 kBq) of ^225^Ac can be delivered with minimal to no bystander toxicities. The pre-targeted interval of 48- 80 hours, depending on mouse strain, was selected for both biodistribution, therapy, and toxicity studies. Proteus clearance studies, in which whole body clearance of hapten in absence of antigen, was performed using [^111^In]In-Pr as a surrogate for [^225^Ac]Ac-Pr. Clearance was rapid, ∼90-94% excreted into the urine by 6 h post-injection (hpi) of tracer,^11^ with no retention in any of the normal organs or tissues.

### 2-step SADA-based alpha-PRIT Dosimetry

The CRC cell line SW1222 (with a GPA33 antigen density of 151,904/cell by quantum bead) was chosen for initial studies. An IV dose of 10 mg/kg GPA33-SADA (step #1) using a 48 h pretargeting interval followed by an IV dose of 37kBq ^225^Ac isotope (1 μCi carried on 700 pmole of proteus) produced an estimated radiation absorbed dose to subcutaneous (sq) tumor, blood, and kidney of 30.88 Gy/μCi, 0.1Gy/μCi, and 2.56 Gy/μCi respectively (TI: 309 for blood; 12 for kidney) (Table 2).

**Table 1:**
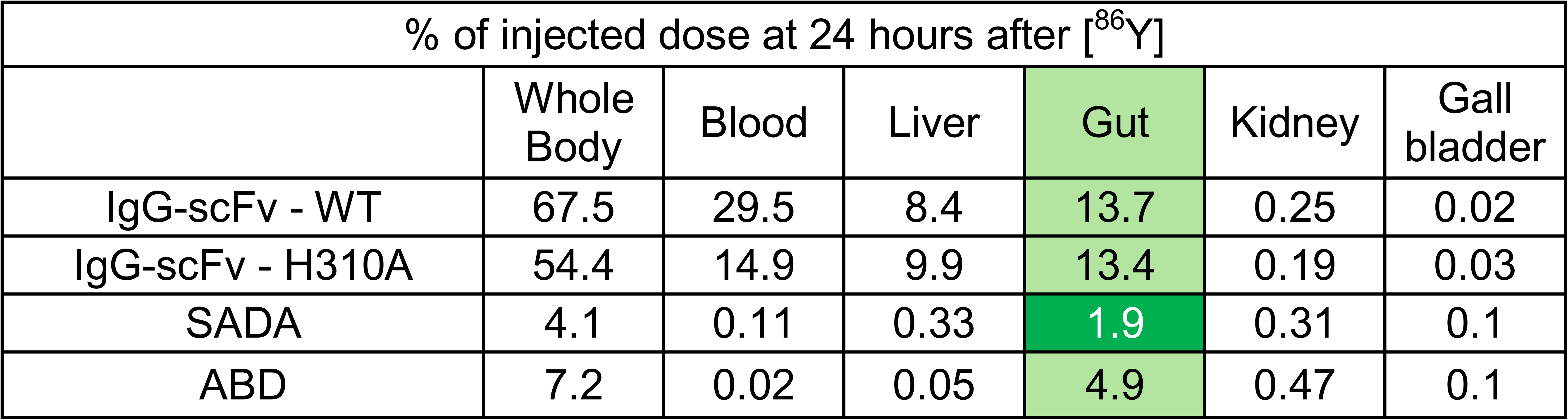
24-hour uptake in cynomolgus NHPs after injection of [^86^Y]Y-ABD combined with different antibody pairs at a 24 h pretargeting interval. Gut uptake was notably lower with SADA BsAb.

| % of injected dose at 24 hours after [<sup>86</sup>Y] |  |  |  |  |  |  |
|---|---|---|---|---|---|---|
|  | Whole Body | Blood | Liver | Gut | Kidney | Gall bladder |
| IgG-scFv - WT | 67.5 | 29.5 | 8.4 | 13.7 | 0.25 | 0.02 |
| IgG-scFv - H310A | 54.4 | 14.9 | 9.9 | 13.4 | 0.19 | 0.03 |
| SADA | 4.1 | 0.11 | 0.33 | 1.9 | 0.31 | 0.1 |
| ABD | 7.2 | 0.02 | 0.05 | 4.9 | 0.47 | 0.1 |

**Table 2:**
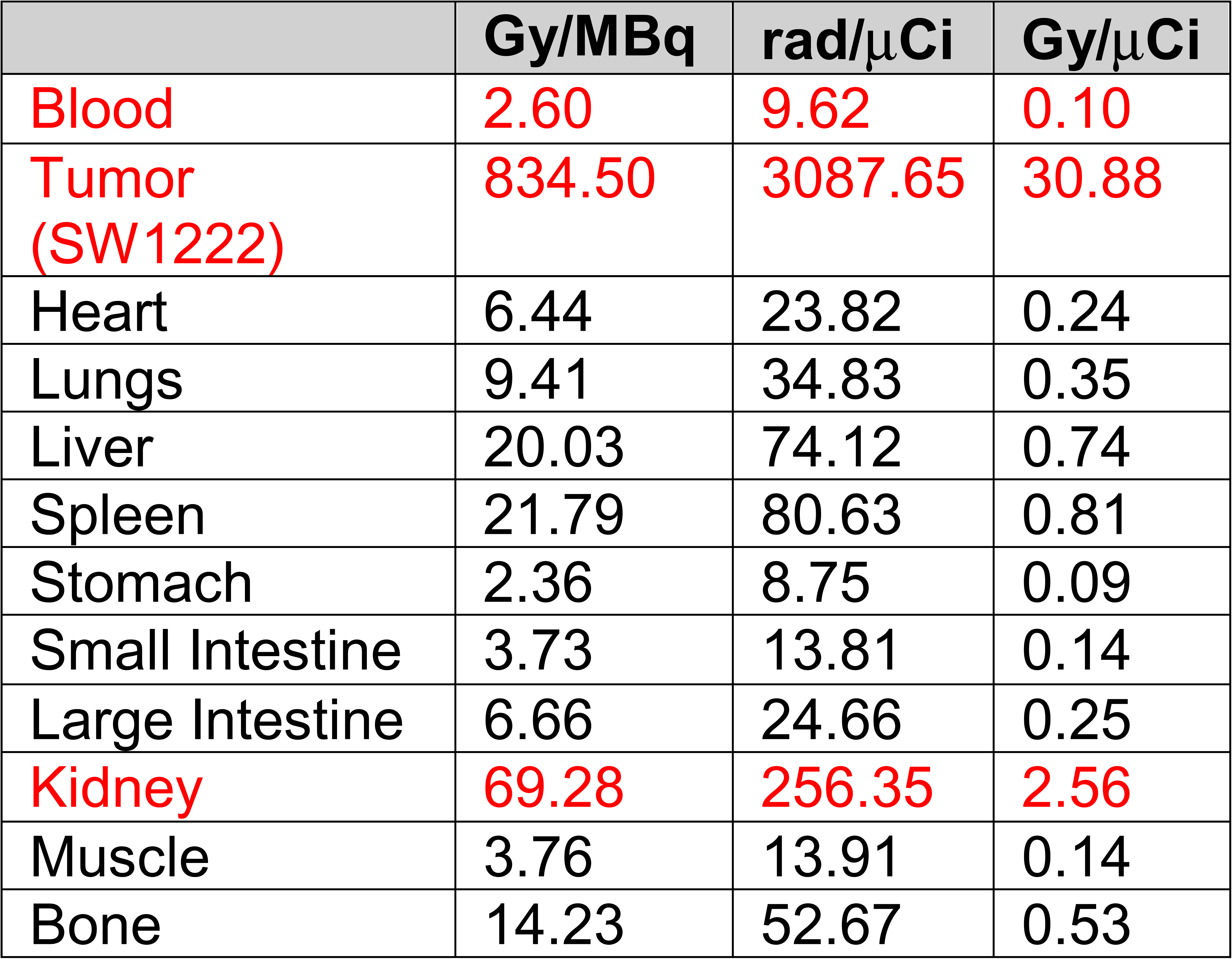
Dosimetry for 37kBq of two-step SADA [225Ac]Ac-Pr in SW1222 murine model.

|  | **Gy/MBq** | **rad/μCi** | **Gy/μCi** |
|---|---|---|---|
| Blood | 2.60 | 9.62 | 0.10 |
| Tumor (SW1222) | 834.50 | 3087.65 | 30.88 |
| Heart | 6.44 | 23.82 | 0.24 |
| Lungs | 9.41 | 34.83 | 0.35 |
| Liver | 20.03 | 74.12 | 0.74 |
| Spleen | 21.79 | 80.63 | 0.81 |
| Stomach | 2.36 | 8.75 | 0.09 |
| Small Intestine | 3.73 | 13.81 | 0.14 |
| Large Intestine | 6.66 | 24.66 | 0.25 |
| Kidney | 69.28 | 256.35 | 2.56 |
| Muscle | 3.76 | 13.91 | 0.14 |
| Bone | 14.23 | 52.67 | 0.53 |

### SADA Can Reduce On-target Off-tumor Uptake in Nonhuman Primates

GPA33-SADA binds to GPA33 in cynomolgus primates. With a pretargeting interval of 24 hours, GPA33 IgG-scFv (0.2 mg/kg IV) at 24 hours post [^86^Y]Y-aminobenzylDOTA ([^86^Y]Y-ABD, 9 nmole) demonstrated expected blood-pool retention typical of IgG-based antibodies and gastrointestinal uptake consistent with prior clinical huA33 immunoPET studies.^12,13^ This on-target off-tumor gut uptake using IgG based constructs did not improve even after introducing a H310A mutation to remove binding to the IgG neonatal receptor FcR(n). In contrast, GPA33-SADA (0.2 mg/kg IV) showed substantially reduced gastrointestinal uptake from 13.4% to 1.9%, even lower than the gut signal observed with radioligand alone (4.9%)(Figure 2, Table 1). Additionally, serial monkey anti-human antibody measurements remained negative over 4 months of follow-up, supporting low immunogenicity of SADA constructs as previously reported.^14^

**Figure 2:**
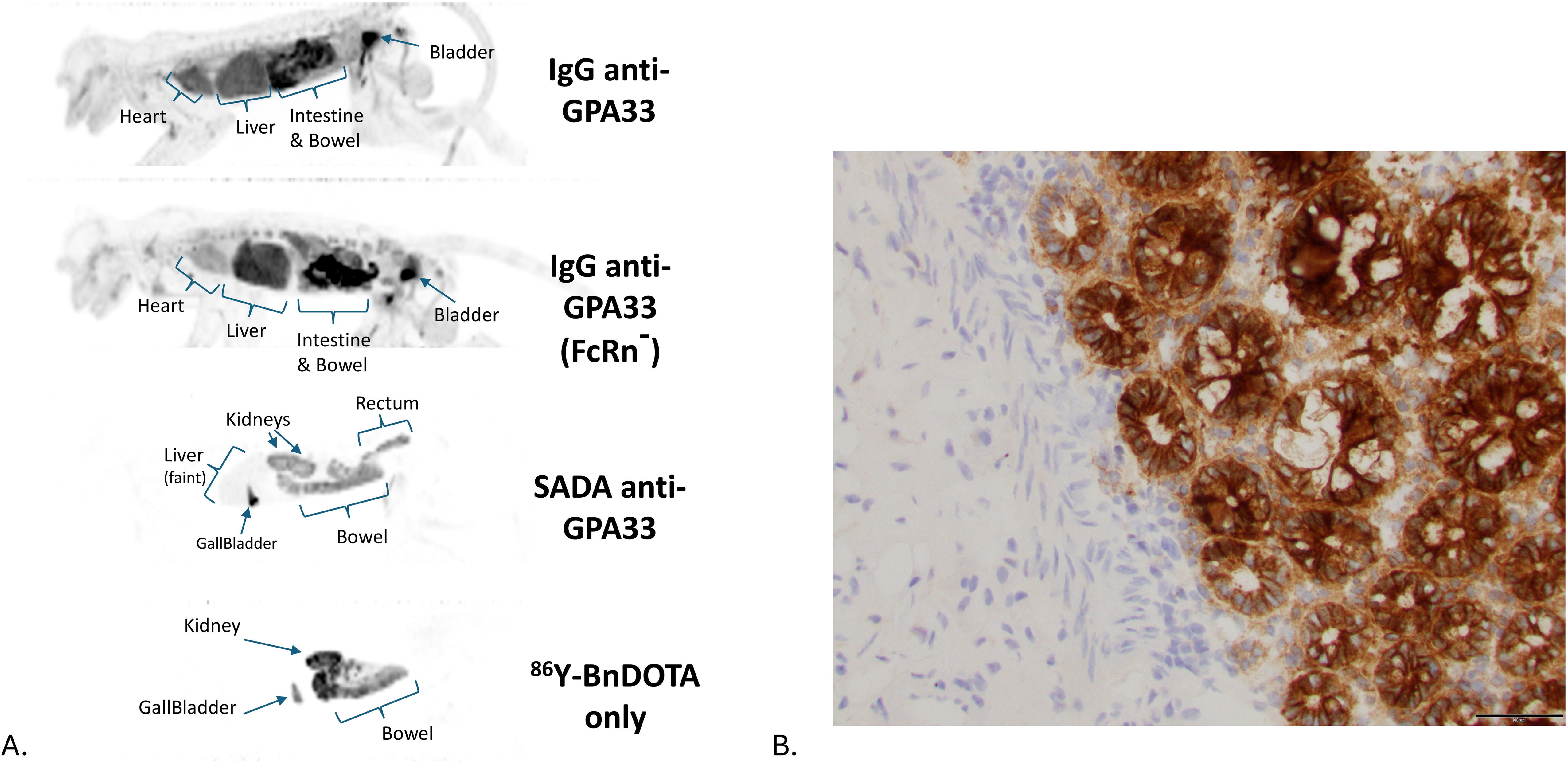
(A) Pretargeting with anti-GPA33 SADA in cynomolgus NHPs leads to significantly reduced gut uptake compared to IgG-scFv BsAb formats. PET imaging was performed 24 hours after [86Y]Y-ABD. BsAb were administered 24 hours prior to [86Y]Y-ABD. **(B)** Immunohisto-chemistry of A33 antigen staining (brown) strong and uniform in colon of cynomolgus monkeys.

At a higher GPA33-SADA dose (10 mg/kg), findings were reproducible across two NHPs, with good tolerability including after repeat dosing in the same animal. Image- derived PK of [^86^Y]Y-DOTA were similar between experiments, with rapid clearance of GPA33-SADA from the blood and most organs within 48-hours and excellent radiohapten clearance by 48-72 hpi. Low-level focal intestinal uptake persisted through 48 hours, consistent with GPA33-specific binding, but remained 3-4 fold lower than previously observed with non-SADA IgG-based constructs. Total activity within the imaging field was approximately 1.5% of the administered dose at 24 hours (5% decay- corrected), while urine HPLC demonstrated only unmetabolized [^86^Y]Y-DOTA and blood/plasma measurements showed minimal residual circulating activity (∼3-fold background)(Figure S1-2).

### Comparing 3-step IgG[L]-scFv-based and 2-step SADA-based alpha PRIT

A subtherapeutic regimen consisting of a single dose of ^225^Ac (74 kBq, 2 μCi) was chosen to compare 3-step versus 2-step alpha PRIT against SW1222 CRC xenografts in nude mice. Median survival (MS; in days after SADA-PRIT treatment start) was 21 d for no treatment and 21 d for treatment with [^225^Ac]Ac-Pr only. In contrast, MS was 58 d for 3-step IgG-scFv BsAb, and 68 d for SADA-BsAb (Log-rank (Mantel Cox); P <0.0001). With the 3-step IgG[L]-scFv BsAb PRIT, 7/8 tumors reached 500 mm^3^ within 75 d, with 0/8 surviving 140 d. In SADA-PRIT, 21/25 reached 500 mm3 within 96 d, with 2/25 surviving 140 d with one mouse showing histologic complete response (CR) (Figure S3).

### Dose Escalation in SW1222 Xenografted Mice

The curative potential of 2 doses versus single dose of alpha-PRIT against sq SW1222 flank tumors in nude mice was compared. Complete tumor volumetric responses were only observed in groups receiving two doses of SADA + ^225^Ac 37Bq+37kBq [^225^Ac]Ac-Pr (2/5), 37kBq+74kBq [^225^Ac]Ac-Pr (1/5) and 74kBq+74kBq [^225^Ac]Ac-Pr) (4/5). MS was 21 d, 21 d, 28 d, 49 d, 78 d, 112 d, and 112 d (p <0.0001) for SADA only, [^225^Ac]Ac-Pr only, irrelevant SADA (GD2-SADA) control group, and GPA33-SADA groups receiving 37kBq, 74kBq, 37+37kBq, and 37+74kBq respectively. MS was not reached in the 74kBq+74kBq [^225^Ac]Ac-Pr group (i.e., >180 d). Two dose regimens were superior to single dose treatment (Figure 3). These results were confirmed in a larger cohort of mice (n=10) (Figure S11). There was no hematological or biochemical evidence of myelotoxicity up to 12 months post treatment (Figure S6). Minor elevations of blood urea nitrogen (BUN) with normal creatinine (Cr) were seen in some mice beginning at 4 months post treatment with higher doses of [^225^Ac]Ac-Pr (Figure S6D, S6E).

**Figure 3:**
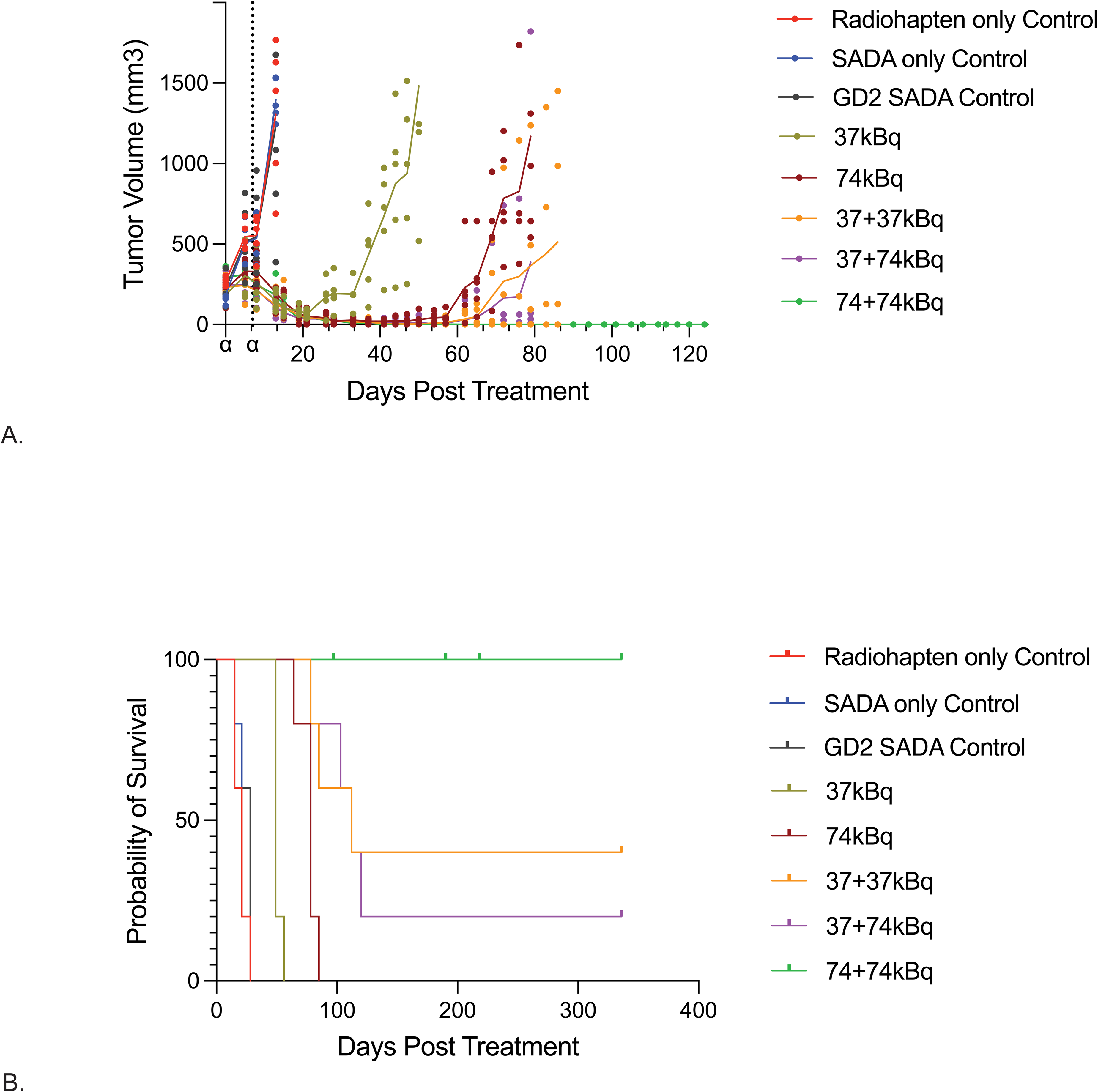
Dose escalation of α-SADA-PRIT with [^225^Ac]Ac-Pr in SW1222. **(A)** Tumor response curves with complete tumor volume response seen in all two dose groups prior to recurrence in all but the 74+74kBq treatment group. **(B)** Median survival was 21 d, 21 d, 28 d, 49 d, 78 d, 112 d, and 112 d (p <0.0001) for SADA only, [^225^Ac]Ac-Pr only, and irrelevant (GD2-SADA) control groups, and GPA33-SADA groups receiving 37kBq, 74kBq, 37+37kBq, and 37+74kBq respectively. MS was not reached for the 74kBq+74kBq (1.7 MBq/kg) group (i.e., >180 d).

### Dose Escalation in Non-Tumor Bearing Mice

A dose escalation study with identical doses was performed in non-tumor bearing mice to ascertain toxicity in the absence of tumor. No hematological evidence of myelotoxicity was observed up to 6 months post injection. Again, some mice, had evidence of BUN elevation beginning at 16 weeks (Figure S7).

### Efficacy Against CRC Xenografts with Diverse Antigen Densities

To build upon the therapeutic and toxicity results, the single cycle two dose regimen of 74kBq+74kBq (∼18 MBq/kg) was tested against two additional sq CRC xenografts- LS174T, a high expressor of GPA33 (216,461/cell by quantum beads), and LOVO, a low expressor of GPA33 (11,342/cell by quantum beads). The two doses were given 7 days apart. With sq LS174T, MS was 24 d and 24 d in SADA only and [^225^Ac]Ac-Pr only control groups, and 140 d in 74kBq+74kBq [^225^Ac]Ac-Pr group (p <0.0001) (Figure 4A). With sq LOVO, MS was 47 d and 60 d in SADA only and [^225^Ac]Ac-Pr only control groups, and 128d in 74kBq+74kBq [^225^Ac]Ac-Pr group (p <0.0001) (Figure 4C). Rate of tumor growth at recurrence was indolent when compared to the rapid initial tumor growth in nonresponding groups (Figure 4B, 4D). All treatments were well tolerated across all groups, with transient reversible weight loss (<15%) and minimal hematologic toxicities (Figure S8-9).

**Figure 4:**
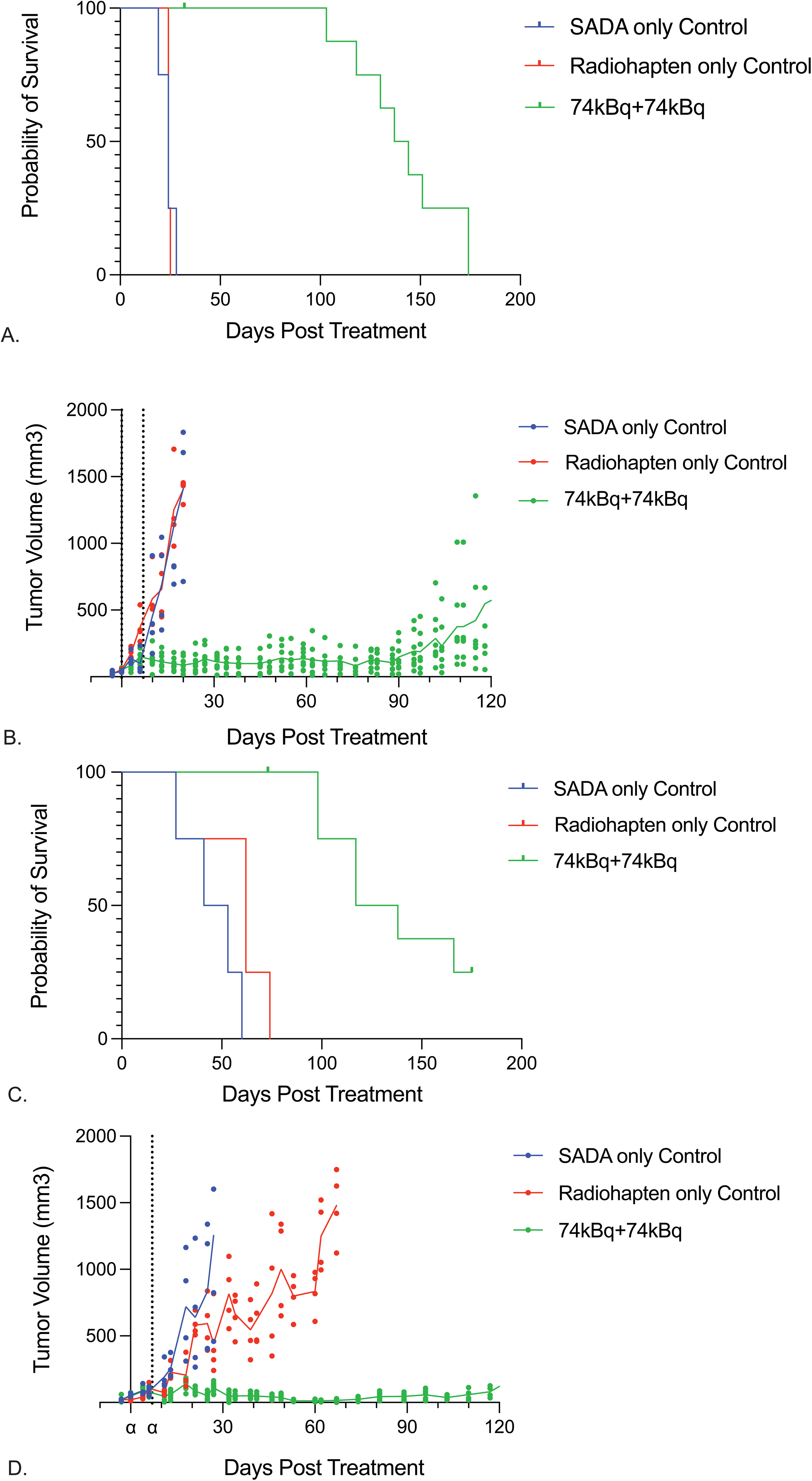
Survival and tumor growth curves in murine models for treatment of sq tumors of CRC cell lines LS174T **(A, B),** a high GPA33 expressor, and LOVO **(C, D)**, a low GPA33 expressor. Treatment dose was 74kBq+74kBq [^225^Ac]Ac-Pr given 7 days apart. In both models, complete tumor response was seen prior to recurrence. Significant survival improvement was observed.

### Efficacy Against Peritoneal Carcinomatosis

In order to test efficacy in a metastatic model, the single cycle two dose (74kBq+74kBq 7 days apart) alpha-SADA PRIT regimen was evaluated in a peritoneal carcinomatosis model, using luciferase transduced SW1222 (SW1222-Lu, antigen density of 89,023/cell by quantum bead) implanted intraperitoneally (ip). Both SADA and [^225^Ac]Ac-Pr were administered IV. MS was 39 d and 40 d in SADA only and [^225^Ac]Ac-Pr only control groups, compared to 134 d in 74kBq+74kBq [^225^Ac]Ac-Pr group (p <0.0001) (Figure 5). There was complete tumor response after 74kBq+74kBq [^225^Ac]Ac-Pr, with 4/10 remaining tumor free at the end point of 140 days. Again, minimal hematological and biochemical toxicities were noted (Figure S10).

**Figure 5:**
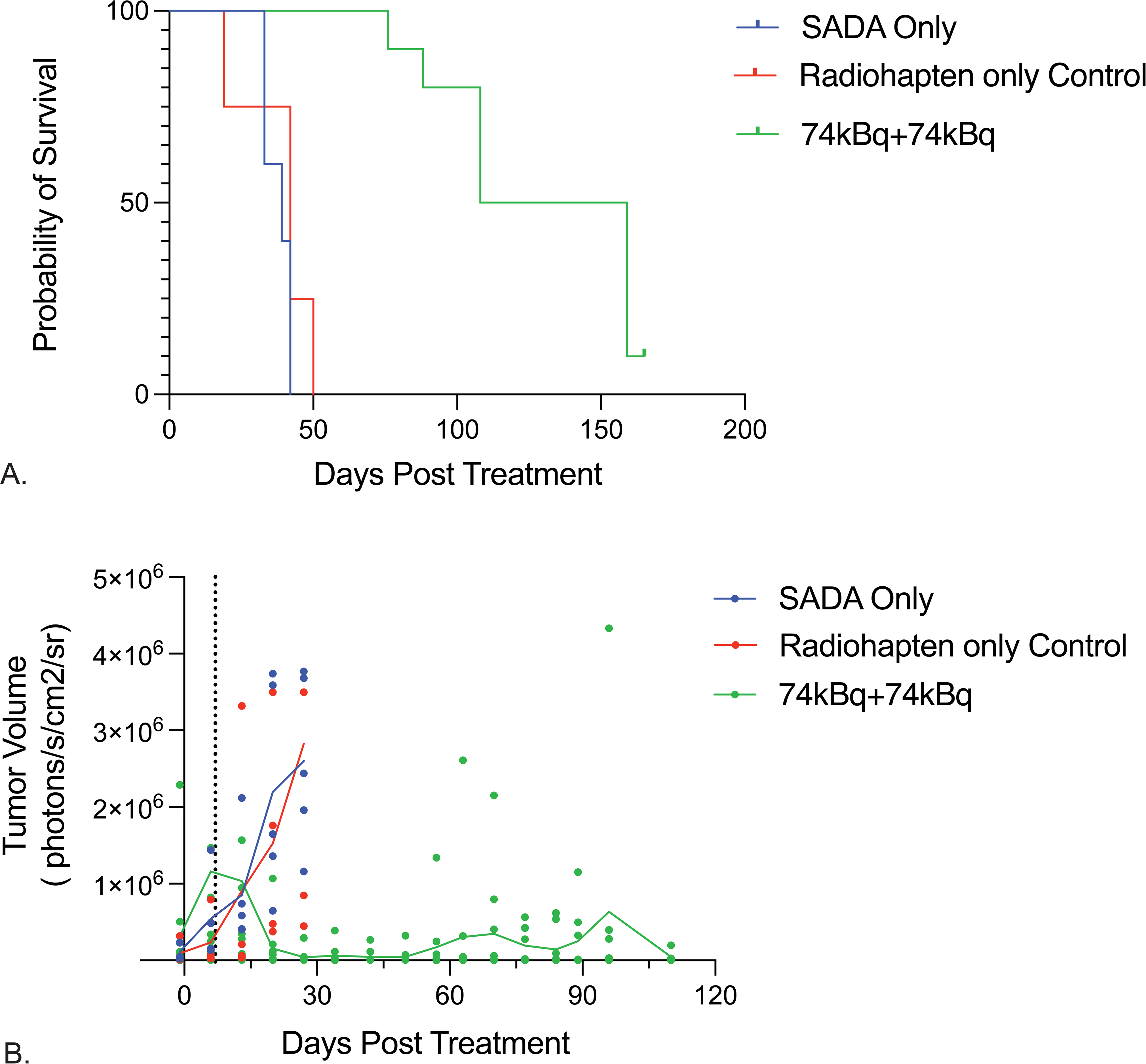
Survival **(A)** and tumor response **(B)** for treatment of an intraperitoneal SW1222 murine model. Complete tumor response was seen in 3/10 mice of the 74kBq+74kBq [^225^Ac]Ac-Pr treatment group. Significantly improved median survival was observed.

### Pathological Evaluation of CRC Xenografts with Diverse Antigen Densities

Histological analysis demonstrated variable treatment responses across tumor models. In the SW1222 sq model, 80% (4/5) mice treated with GPA33-SADA + 74kBq+74kBq [^225^Ac]Ac-Pr were tumor free, or histologically cured, with one showing only a small fibrocollagenous nodule at the implantation site without metastasis (Figure 6A).

**Figure 6:**
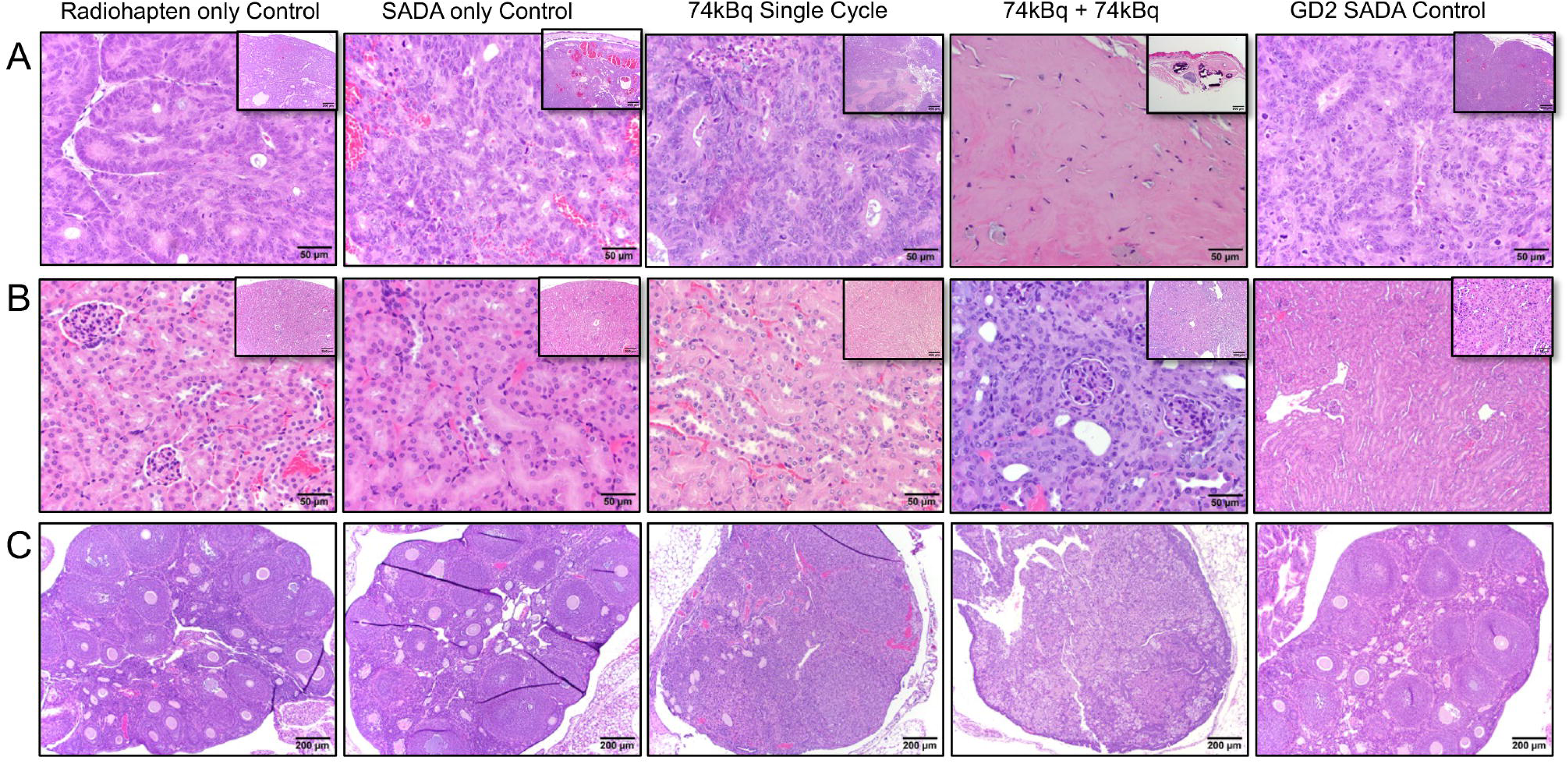
Representative histopathology of tumors, kidneys, and ovaries in athymic nude mice bearing sq SW1222 tumors treated with SADA PRIT. **(A)** Tumor xenografts consistent with adenocarcinoma were observed in mice from the following treatment groups: 74KBq + 74kBq ^225^Ac only (Group 1), SADA only (Group 2), SADA with single dose 74kBq ^225^Ac (Group 4), and 74kBq + 74kBq ^225^Ac with irrelevant GD2-SADA (Group 8). In contrast, mice treated with SADA + 74kBq + 74kBq ^225^Ac (Group 7) exhibited a fibrocollagenous nodule containing rare, scattered neoplastic cells at the site of tumor cell injection (40x magnification; inset: 10x mag.). **(B)** Renal histology appeared normal in mice from Groups 1, 2, 4, and 8. In contrast, representative kidneys from Group 7 mice showed mild to moderate tubular degeneration and regeneration, tubular atrophy and attenuation, and rare karyomegalic epithelial cells (40x mag.; inset: 10x mag.). **(C)** Ovarian histology was unremarkable in mice from Groups 1, 2, 4, and 8. However, ovaries from Group 7 mice exhibited mild to moderate cortical atrophy with follicular depletion and ceroid pigment accumulation (10x mag).

LS174T-engrafted mice had microscopic residual carcinoma at the injection site at 140 days, with one developing distant metastasis (Figure S4A). In LOVO tumors, 4/9 mice were tumor-free and 5/9 had carcinomas with necrosis and mineralization (Figure S5A). In the ip SW1222 model, 2/5 mice were cured and 3/5 had small recurrent adenocarcinomas.

Histologic evidence of dose-dependent renal tubular injury was seen in the SW1222 (Figure 6B; Table S3), LS174T (Figure S4B; Table S5), and LOVO (Figure S5B; Table S7) sq models treated with GPA33-SADA + [^225^Ac]Ac-Pr, ranging from minimal to marked tubulointerstitial pathology. Control kidneys (SADA alone or [^225^Ac]Ac-Pr alone) were normal or showed only mild background changes. All ip SW1222 mice developed histologic evidence of renal radiotoxicity, absent in controls (Table S9).

Ovarian histopathology exhibited mild-to-marked cortical atrophy and follicular degeneration in mice receiving two dose treatment in the SW1222 dose escalation, which was notably absent in single dose (SADA + 74kBq [^225^Ac]Ac-Pr ) or control groups (Figure 6C; Table S4). Similar ovarian findings were detected in sq LS174T (Figure S4C; Table S6), sq LOVO (Figure S5C; Table S8) and ip SW1222 (Table S10) models treated with 74 + 74 kBq while controls were unremarkable.

No radiotoxic lesions were identified in the bone marrow, spleen, liver, urinary bladder, or peripheral lymph nodes. Minor microscopic findings across cohorts were interpreted as physiological, reactive, or incidental age- or strain-related changes.^15^ A single sq SW1222 bearing treatment mouse developed spontaneous lymphoma, a finding within the institutional incidence of 10% in both experimental and littermate controls.

In non-tumor bearing mice, toxicity profiling was consistent with tumor bearing counterparts. At study endpoint of 120 days, all mice treated with two 74 kBq doses showed minimal to marked renal tubular lesions, while single-dose kidneys were largely normal (Table S1). Ovarian atrophy occurred in multi-dose groups but not controls (Table S2). Despite these histologic changes, body weights of mice at necropsy remained stable over months without clinical toxicities before necropsy.

## DISCUSSION

This study demonstrates that two-step GPA33-SADA pretargeted alpha- radioimmunotherapy can deliver therapeutically active ^225^Ac to colorectal cancer while substantially reducing exposure to normal GPA33-expressing gastrointestinal tissue. Across subcutaneous and intraperitoneal CRC xenograft models spanning a broad range of GPA33 expression, a single treatment cycle consisting of two doses of [^225^Ac]Ac-Pr produced deep and durable tumor responses, prolonged survival, and histologically confirmed tumor-free animals in several models. These antitumor effects were achieved without clinically apparent acute hematologic or biochemical toxicity, although repeat treatment produced delayed renal tubular and ovarian histopathologic injury. Together, these findings support the potential of SADA-based pretargeting to widen the therapeutic window of alpha-particle therapy while identifying kidney and gonadal exposure as important constraints for further optimization.

Radiopharmaceutical therapy (RPT) has advanced rapidly with improvements in chelation chemistry, reliable isotope sourcing, and ligand engineering. Clinically established examples include SIR-Spheres ^90^Y resin for hepatic tumors, Xofigo (^223^Ra- dichloride) and ^177^Lu- and ^225^Ac-PSMA-617 for metastatic castrate-resistant prostate cancer^16^, and Lutathera (^177^Lu-DOTATATE) for neuroendocrine tumors^17^. A central challenge, however, remains the delivery of sufficiently high tumor absorbed doses without exceeding normal-tissue tolerance. Tumor response in RPT has been associated with absorbed dose ^18,19^ in line with the clinical experience of external beam radiation, where >50 Gy doses are generally required for consequential tumor responses.^5^ Yet, as highlighted by the Radiopharmaceutical Therapy-Tissue Effects in the Clinic (RPT-TEC) group, dose constraints in RPT are often extrapolated from external-beam organ limits rather than observed radiotoxicities, perpetuating underdosing, while narrow therapeutic windows remain a major hurdle.^20^

GPA33 illustrates an additional challenge because the antigen is highly expressed not only by CRC but also by normal intestinal epithelium. Conventional GPA33-targeted IgG approaches therefore face both systemic exposure from prolonged antibody circulation and on-target, off-tumor intestinal binding.^4,21–24^ The SADA platform addresses both limitations through size-dependent pharmacokinetics: tumor-bound tetramers are retained, whereas unbound molecules dissociate into smaller species that undergo rapid clearance before administration of the radioactive hapten. In cynomolgus nonhuman primates, GPA33-SADA markedly reduced gastrointestinal uptake compared with IgG-scFv constructs, while retaining rapid systemic clearance (**Table 1, Figure S1**). This behavior was reproduced at a 10 mg/kg SADA dose and after repeat administration. Importantly, these studies were performed in animals without a GPA33- positive tumor sink, providing a stringent assessment of normal-tissue binding. These observations support the possibility that SADA pharmacokinetics can mitigate the gastrointestinal exposure that has constrained previous GPA33-directed approaches.

The resulting biodistribution translated into a favorable therapeutic index in the SW1222 model. Dosimetry following GPA33-SADA pretargeting yielded tumor-to-blood and tumor-to-kidney absorbed-dose ratios of 309 and 12, respectively (**Table 2**). This therapeutic separation was accompanied by substantial antitumor activity. Two-dose treatment was superior to single-dose treatment in SW1222 tumors, and the 74 kBq + 74 kBq regimen produced durable responses in SW1222, LS174T, and LOVO subcutaneous xenografts despite marked differences in GPA33 antigen density (**Figure 3-4**). Activity was also maintained in the intraperitoneal SW1222 model, where treatment prolonged median survival and produced tumor-free survivors (**Figure 5)**.

Histopathologic assessment further demonstrated complete elimination of viable tumor in subsets of treated animals after extended follow-up of 6 months, which is beyond typical endpoints used in most preclinical RPT studies.^25^ Collectively, these findings suggest that the combination of rapid normal-tissue clearance and high-linear-energy- transfer alpha radiation can generate meaningful therapeutic activity even when antigen expression is relatively low or disease is distributed within the peritoneal cavity.

These results also illustrate the potential advantages of SADA relative to earlier three- step IgG-based PRIT strategies. Conventional PRIT can improve tumor-to-normal- tissue ratios but requires a separate clearing agent and introduces additional pharmacokinetic and operational complexity.13,26-32 SADA reduces this process to two administrations while retaining the fundamental advantage of separating tumor targeting from radionuclide delivery. The ability to modulate antibody dose, pretargeting interval, radiohapten activity, and treatment fractionation provides several opportunities for further optimization. Whether the simplified pharmacology of SADA translates into greater reproducibility or a wider therapeutic window in humans will require prospective clinical evaluation.

These results also illustrate the potential advantages of SADA relative to earlier three- step IgG-based PRIT strategies. ^26^ Conventional PRIT can improve tumor-to-normal- tissue ratios but requires a separate clearing agent and introduces additional pharmacokinetic and operational complexity.^13,27–32^ SADA reduces this process to two administrations while retaining the fundamental advantage of separating tumor targeting from radionuclide delivery. The ability to modulate antibody dose, pretargeting interval, radiohapten activity, and treatment fractionation provides several opportunities for further optimization. Whether the simplified pharmacology of SADA translates into greater reproducibility or a wider therapeutic window in humans will require prospective clinical evaluation.

The therapeutic activity observed here must be considered together with the delayed histopathologic findings. Repeat ^225^Ac-SADA-PRIT produced dose-dependent renal tubular injury across tumor-bearing models and in non-tumor-bearing mice, despite largely preserved serum renal indices and the absence of overt clinical toxicity. Ovarian cortical atrophy and follicular depletion were also observed predominantly after two- dose treatment. These findings are consistent with the kidney serving as an important route of clearance for ^225^Ac-associated activity and its radioactive daughters.^11,33–34^ Ovarian cortical atrophy and follicular loss observed in some mice are similarly consistent with nonspecific cross-irradiation by nearby radioactive tissues.^9,28,35^ Though not life-threatening, given the radiosensitivity of gonads,^36^ even 200 cGy from proximity to the kidney or misplaced ^225^Ac decay daughters can cause damage.^37^ Importantly, there were no dose-limiting toxicities involving marrow, liver, or nerve tissue, and histologic evaluation revealed only renal and ovarian changes without evidence of life-threatening organ damage (**Figure 6, Figure S4-5, Table S1-S10**). Collectively, these findings support a favorable preclinical therapeutic index while highlighting renal and gonadal exposure as important considerations for clinical translation. Strategies to reduce renal radiation exposure and characterize gonadal dosimetry will be essential to further optimize the therapeutic window of this alpha-PRIT platform.

Several limitations should inform interpretation and clinical translation. Therapeutic efficacy was evaluated in immunodeficient xenograft models, which do not fully reproduce the tumor microenvironment, antigen distribution, or systemic physiology of patients with metastatic CRC. Species-specific differences in pharmacokinetics, radiation sensitivity, and ^225Ac daughter redistribution may further limit extrapolation of murine dosimetry and toxicity findings. Although the NHP studies provide important evidence of reduced gastrointestinal uptake in a species with relevant GPA33 expression, they were not designed to establish therapeutic safety. Additionally, the current data do not define the minimum GPA33 expression required for response or the optimal relationship among targeting-agent dose, radionuclide activity, and treatment interval. These questions will be important for establishing clinically appropriate treatment regimens and identifying patients most likely to benefit.

Clinical development should prioritize individualized imaging and dosimetry to characterize tumor targeting and normal-organ exposure before therapeutic administration. Diagnostic radiohaptens, including ^86Y-labeled constructs, may provide a means of evaluating biodistribution and informing patient-specific treatment planning. Early-phase studies should establish absorbed-dose relationships in tumor, kidney, gastrointestinal tract, marrow, and gonadal tissues, with particular attention to delayed toxicity and the feasibility of repeat treatment. Further preclinical optimization should address renal protection, antigen-density thresholds, and treatment strategies for heterogeneous disease.

Collectively, these findings demonstrate that SADA-mediated pretargeting can achieve durable antitumor activity while substantially reducing gastrointestinal exposure associated with conventional GPA33-directed targeting. The combination of favorable murine dosimetry, durable responses across heterogeneous CRC models, and reduced intestinal uptake in NHPs provides a compelling rationale for clinical development. More broadly, this approach suggests that pharmacokinetic separation of antigen recognition and radionuclide delivery may expand the therapeutic potential of tumor-associated antigens whose expression in normal tissues has historically limited their clinical utility. Establishing whether this therapeutic separation can be reproduced in humans, while mitigating renal and other delayed toxicities, represents the critical next step toward clinical translation of GPA33-SADA α-PRIT.

## List of Supplementary Materials

Figure S1-S11 Table S1-S10

## Acknowledgements

The authors would like to acknowledge the United States Department of Energy for the provision of Actinium-225. The authors are grateful to Dr. Sara Rinne for thoughtful editorial input and careful review of the manuscript.

## Author Contribution

Conceptualization: NA, SC, AC, MBF, DV, SMC, GK, SML, GMN, SKC, DV, NKC Methodology: NA, NV, SC, SGL, HFG, SEC, RM, EKF, HX, YF, IC, DV, NKC Investigation: NA, NV, SGL, HFG, ALR, JAC, MK, KL, NKC

Visualization: NA, NV, SC, SEC, RM, EKF, ALR, JAC, IC, DV, NKC

Funding acquisition: NA, AC, MBF, DV, SMC, DV, GMN, NKC Project administration: NA, SC, MK, KL, NKC

Supervision: SML, DV, GMN, NKC

Writing – original draft: NA, SC, SEC, RM, NKC

Writing – review & editing: NA, NV, DV, SMC, GK, SML, GMN, NKC

## Data and Materials Availability

All data are available in the main text or the supplementary materials.

