## Supplemental Files for "Actinium-225 Pretargeted Radioimmunotherapy against Colorectal Cancer using Self-Assembling and Disassembling Antibodies"

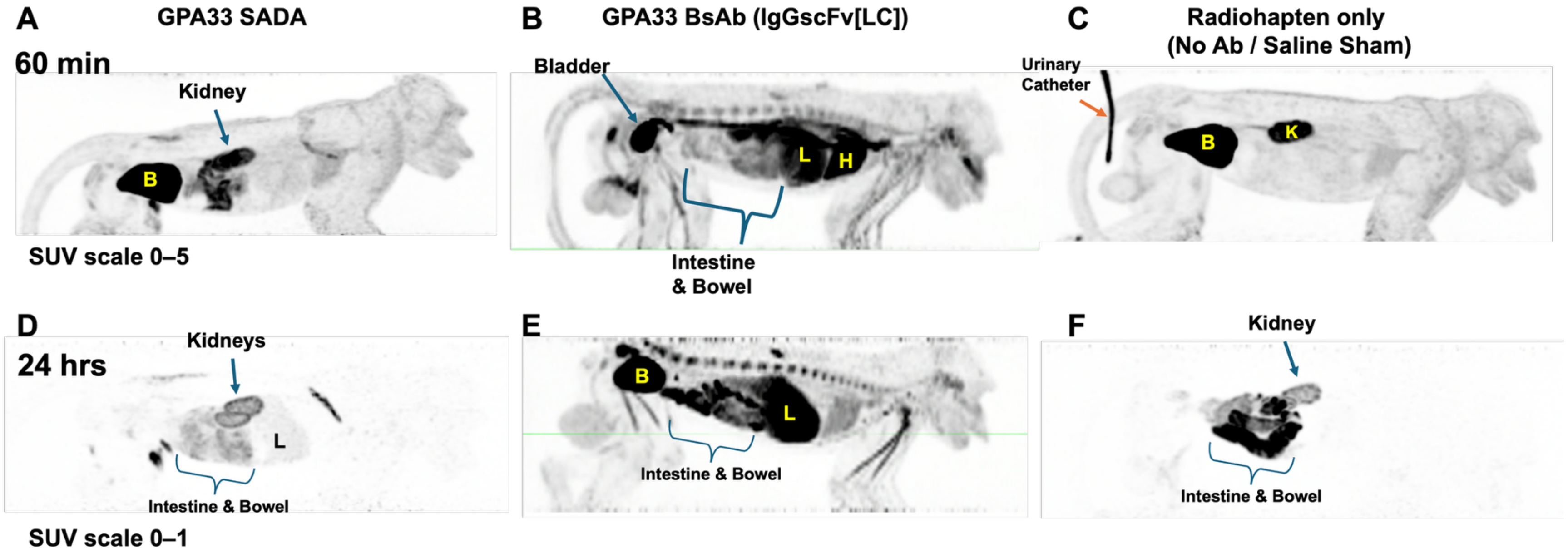


**Figure S1:** $^{86}$Y PET imaging at 1 hpi and 24 hpi pretargeted with 10mg/kg GPA33 SADA **(A, D)** or 0.25mg/kg bispecific IgGscFv[LC] antibody **(B,E)**, or no antibody **(C,F).**

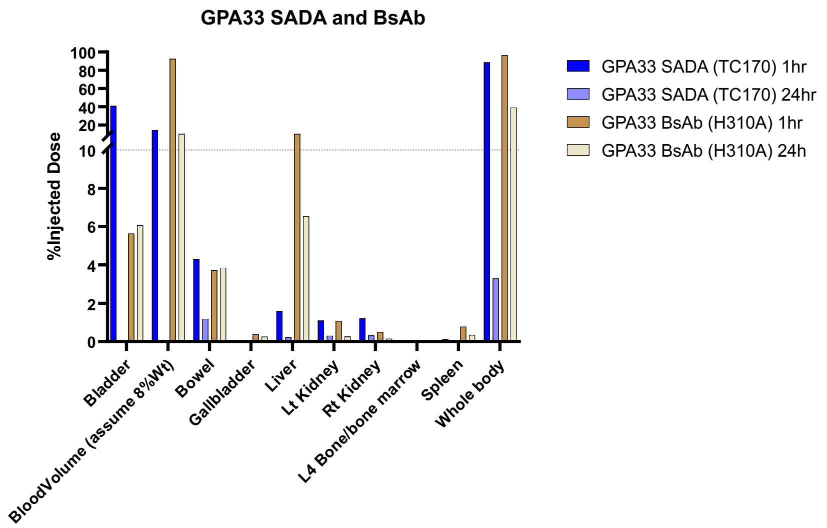


A.

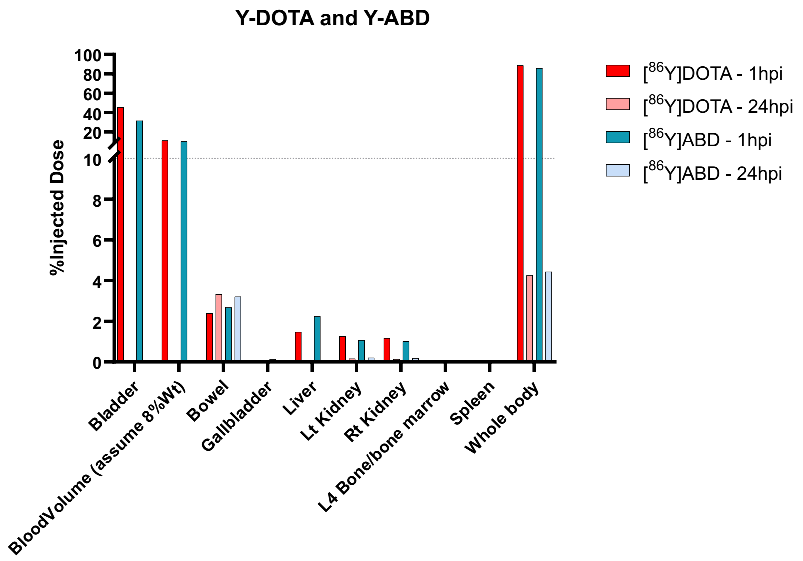


B.

**Figure S2:** Image based biodistribution in NHP. **(A)** PET/CT-derived biodistribution of GPA33-SADA compared with the IgG H310A variant demonstrated low gastrointestinal accumulation with <4% uptake in the bowel. **(B)** Biodistribution of [86Y]Y-DOTA compared with [86Y]Y-ABD at 1 hpi and 24 hpi demonstrated similarly low bowel uptake (~2%), with rapid clearance of radioligand from non-target tissues.

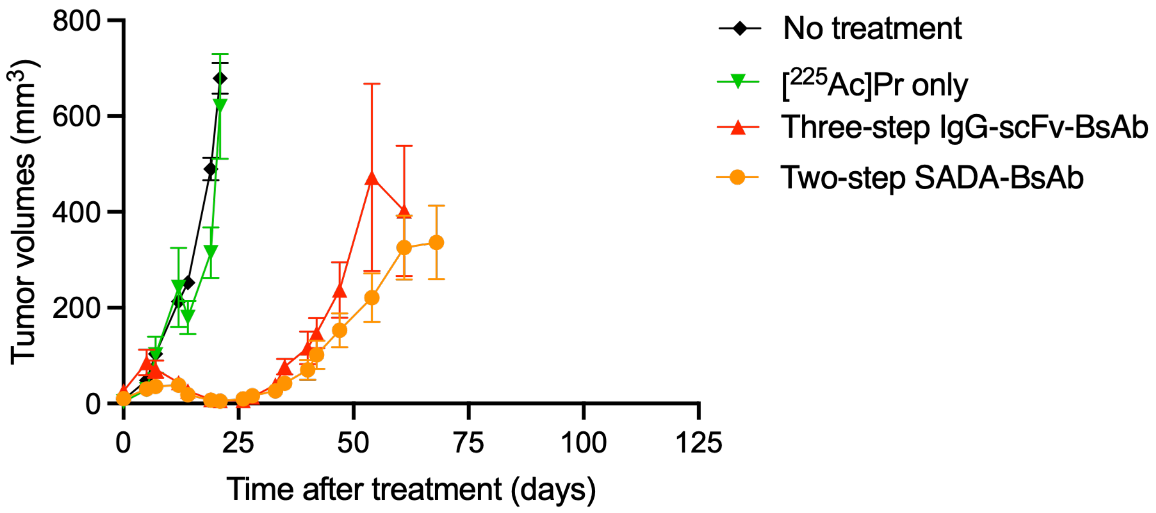


**Figure S3:** Tumor response curves of 3-step IgG-scFv BsAb compared to 2-step SADA-BsAb. Data is presented as average ± SEM. For each group, data is shown until MS was reached.

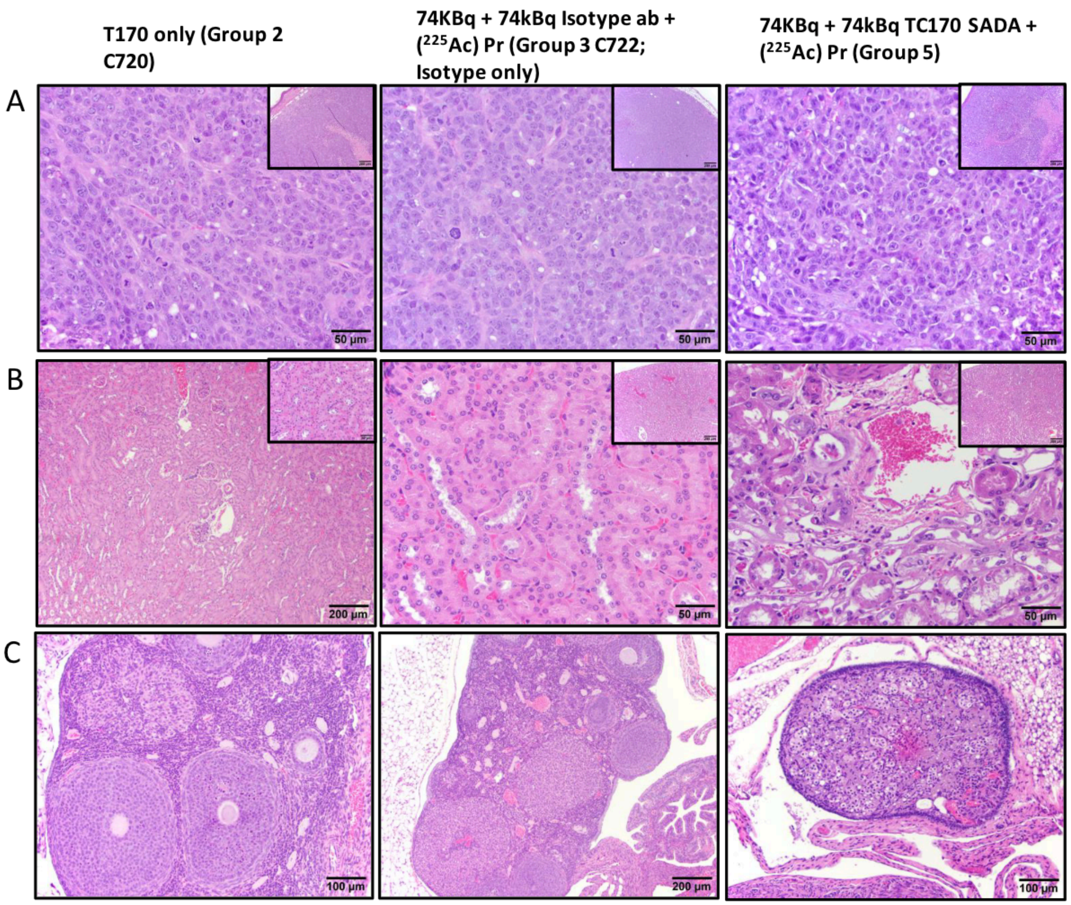


**Figure S4:** Representative histopathology of tumors, kidneys, and ovaries in athymic nude mice bearing sq LS174T tumors treated with SADA + [$^{225}$Ac]Ac-Pr PRIT. **(A)** Tumor xenografts consistent with carcinoma were observed in mice from the following treatment groups: SADA only, 74KBq + 74kBq [$^{225}$Ac]Ac-Pr only, and treatment 74KBq + 74kBq SADA + [$^{225}$Ac]Ac-Pr (40x magnification; inset: 10x mag.). **(B)** Representative histology of normal murine kidneys from Groups 2 and 3. In contrast, kidneys from Group 5 mice exhibited mild to moderate tubular attenuation and degeneration, with occasional karyomegalic epithelial cells (40x mag.; inset: 10x mag.). **(C)** Ovarian histology was unremarkable in mice from Groups 2 and 3. However, ovaries from Group 5 mice showed moderate to marked cortical atrophy accompanied by follicular loss (10x mag.).

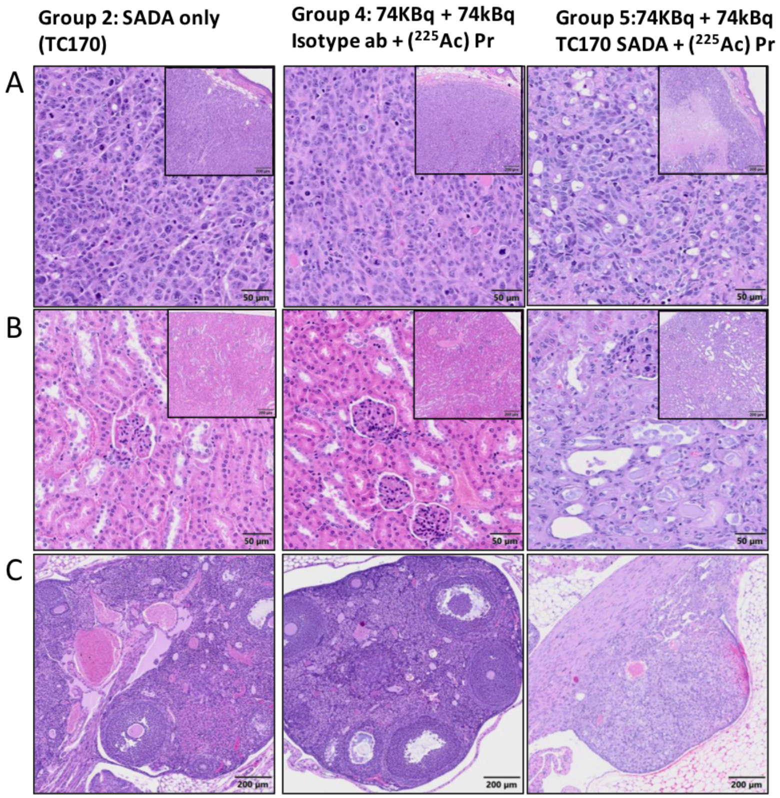


**Figure S5:** Representative histopathology of tumors, kidneys, and ovaries in athymic nude mice bearing sq LOVO tumors treated with SADA + [$^{225}$Ac]Ac-Pr PRIT. **(A)** Tumor xenografts consistent with carcinoma were observed in mice from the following treatment groups: SADA only, 74KBq + 74kBq [$^{225}$Ac]Ac-Pr only, and treatment 74KBq + 74kBq SADA + [$^{225}$Ac]Ac-Pr. The carcinoma from a representative Group 5 mouse exhibited prominent intratumoral necrosis (20x magnification; inset: 10x mag.). **(B)** Representative histology of normal murine kidneys from Groups 2 and 4. In contrast, kidneys from Group 5 mice exhibited mild to moderate tubular attenuation, degeneration, and ectasia with luminal proteinaceous content. Minimal, segmental hyaline glomerulopathy is observed in representative Group 5 kidney (40x mag.; inset: 20x mag.). **(C)** Ovarian histology was unremarkable in mice from Groups 2 and 4. However, ovaries from Group 5 mice showed moderate to marked cortical atrophy accompanied by follicular loss (10-20x mag.).

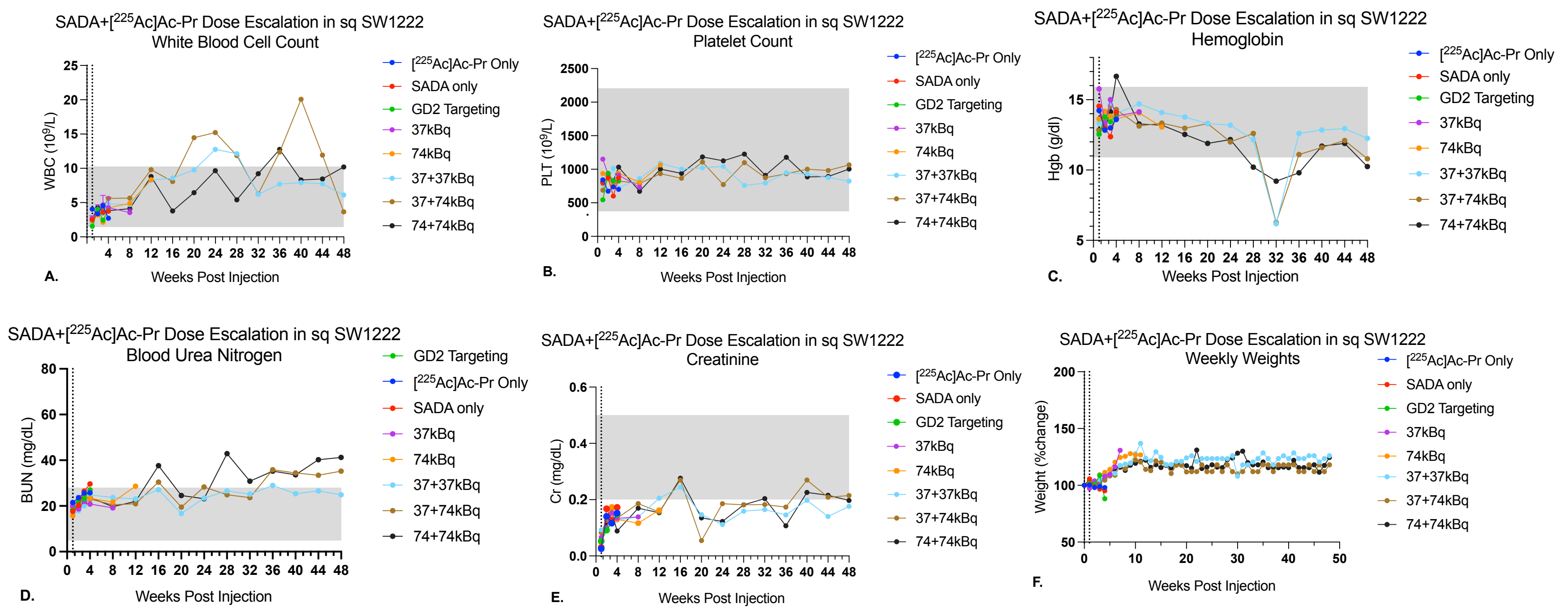


*First dose given at time point 0, second at 7 days, specified by dotted line at x-axis.

**Figure S6:** Representative hematology and renal makers from dose escalation with SADA + [$^{225}$Ac]Ac-Pr. Groups were [$^{225}$Ac]Ac-Pr only, SADA only, irrelevant SADA (GD2-SADA) control group, GPA33-SADA treatment groups receiving 37kBq, 74kBq, 37+37kBq, 37+74kBq, and 74kBq+74kBq. **(A)** White blood cell count (WBC) demonstrating no period of leukopenia for any dose given. Periods of leukocytosis in later months of the study are likely due to infection or disease burden in these older mice. On occasion this correlated to lymphoma, as described in the pathological results. **(B)** Platelet (Plt) demonstrating no period of thrombocytopenia for any dose given. **(C)** Hemoglobin (Hgb) demonstrating anemia in all groups at 32 weeks, possibly correlative with laboratory error. **(D)** Blood Urea Nitrogen (BUN), a maker of renal function, demonstrating elevation in the 74kBq+74kBq treatment group beginning at 16 weeks post injection as well as the other two dose treatment groups beginning at 28 weeks post injection, which correlated with renal tubular pathology seen on histopathological evaluation. **(E)** Creatinine (Cr) demonstrating no change from baseline range throughout the study period. **(F)** Weights, measured as a % change from baseline at week 0, exhibiting increase in size over time of all groups.

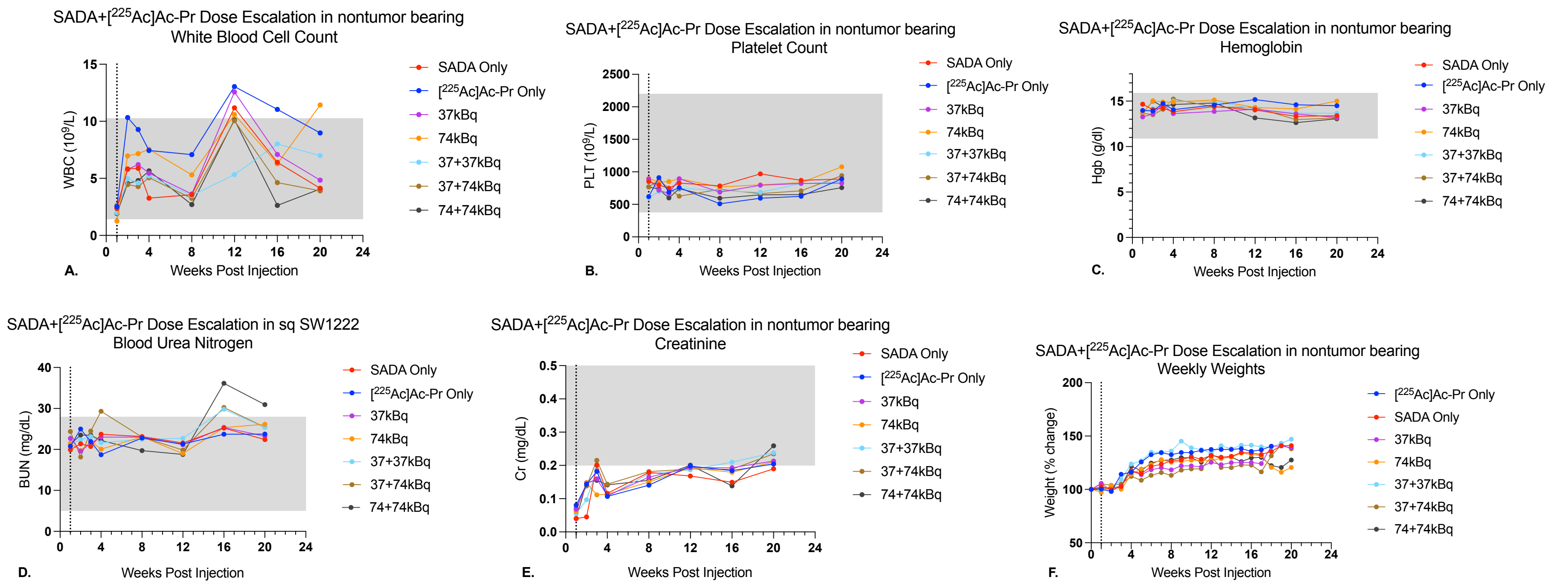


**Figure S7:** Representative hematology and renal makers from dose escalation with SADA + [$^{225}$Ac]Ac-Pr in non tumor bearing mice. Groups were [$^{225}$Ac]Ac-Pr only, SADA only, GPA33-SADA treatment groups receiving 37kBq, 74kBq, 37+37kBq, 37+74kBq, and 74kBq+74kBq. Mice were followed for a period of 20 weeks. **(A)** White blood cell count (WBC) demonstrating no period of leukopenia for any dose given. **(B)** Platelet (Plt) demonstrating no period of thrombocytopenia for any dose given. **(C)** Hemoglobin (Hgb) demonstrating no period of anemia for any dose given. **(D)** Blood Urea Nitrogen (BUN), a maker of renal function, demonstrating a mild elevation in two dose treatment groups beginning at 16 weeks. Again correlating with histopathological evaluation of minimal to moderate tubular injury. **(E)** Creatinine (Cr) demonstrating no change from baseline range throughout the study period. **(F)** Weights, measured as a % change from baseline at week 0, exhibiting increase in size over time of all groups.

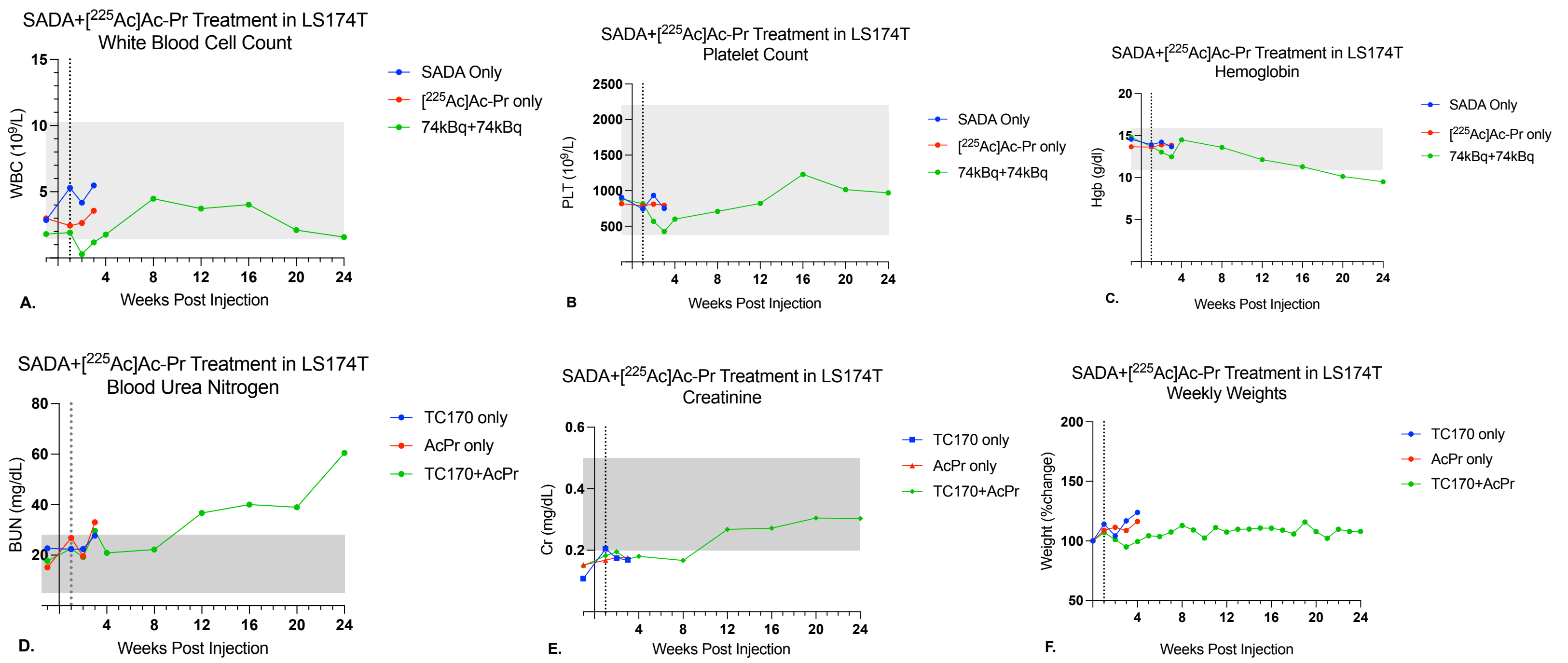


**Figure S8:** Representative hematology and renal markers from treatment of LS174T with SADA + [$^{225}$Ac]Ac-Pr. Groups were [$^{225}$Ac]Ac-Pr only, SADA only, GPA33-SADA treatment receiving 74kBq+74kBq. Mice were followed for a period of 24 weeks. **(A)** White blood cell count (WBC) demonstrating transient leukopenia in the treated group at week 2 post injection. **(B)** Platelet (Plt) demonstrating no period of thrombocytopenia. **(C)** Hemoglobin (Hgb) demonstrating anemia near study endpoint, likely correlating to disease burden effects. **(D)** Blood Urea Nitrogen (BUN), a maker of renal function, demonstrating an elevation in treatment group beginning at 12 weeks, correlating with histopathological findings. **(E)** Creatinine (Cr) demonstrating no change from baseline range throughout the study period. **(F)** Weights, measured as a % change from baseline at week 0, exhibiting stable weight over time in treated mice.

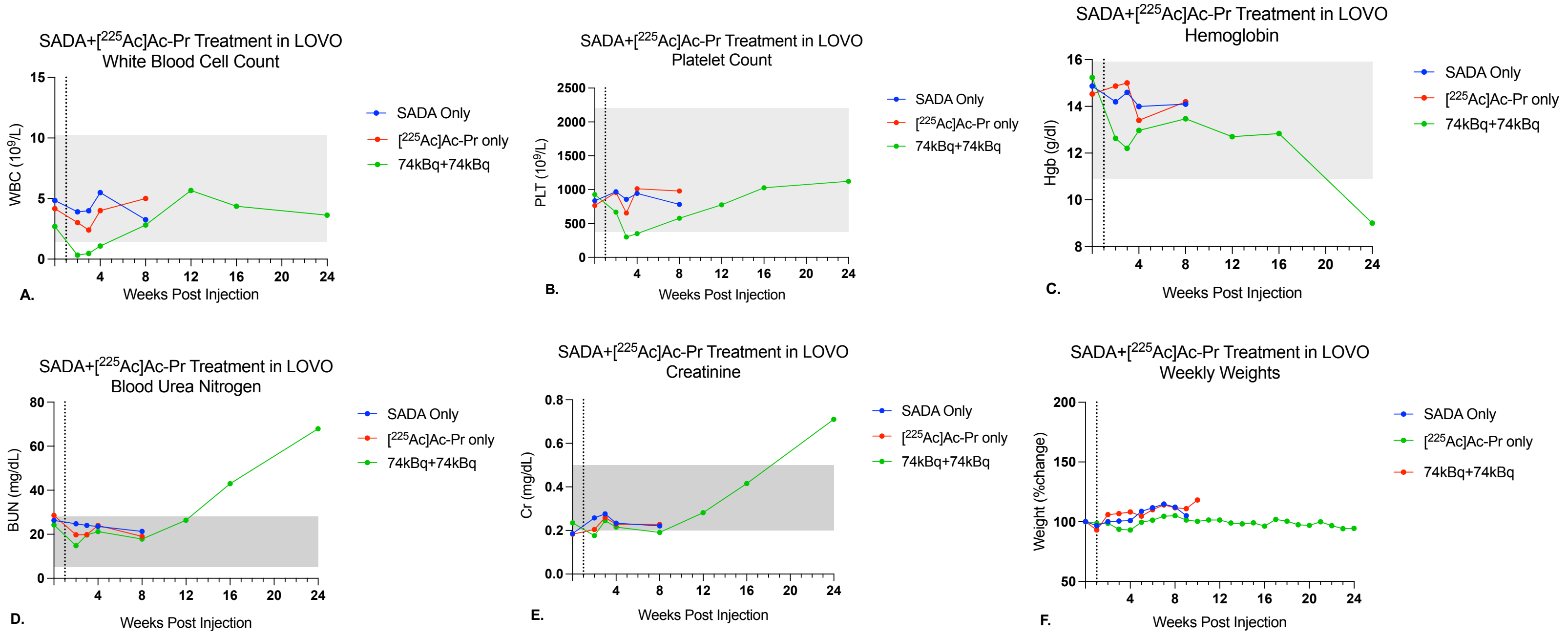


**Figure S9:** Representative hematology and renal markers from treatment of LOVO with SADA + [$^{225}$Ac]Ac-Pr. Groups were [$^{225}$Ac]Ac-Pr only, SADA only, GPA33-SADA treatment receiving 74kBq+74kBq. Mice were followed for a period of 24 weeks. **(A)** White blood cell count (WBC) demonstrating transient leukopenia in the treated group at week 2 post injection. **(B)** Platelet (Plt) demonstrating no period of thrombocytopenia. **(C)** Hemoglobin (Hgb) demonstrating anemia near study endpoint, likely correlating to disease burden effects. **(D)** Blood Urea Nitrogen (BUN), a maker of renal function, demonstrating an elevation in treatment group beginning at 16 weeks, correlating with histopathological evaluation. Steep increase at study endpoint could also correlate with dehydration in the setting of disease burden. **(E)** Creatinine (Cr) demonstrating increase at study endpoint, which is not reflected in any other study group, possibly correlating with dehydration in the setting of disease burden. **(F)** Weights, measured as a % change from baseline at week 0, exhibiting increase in size over time of all groups.

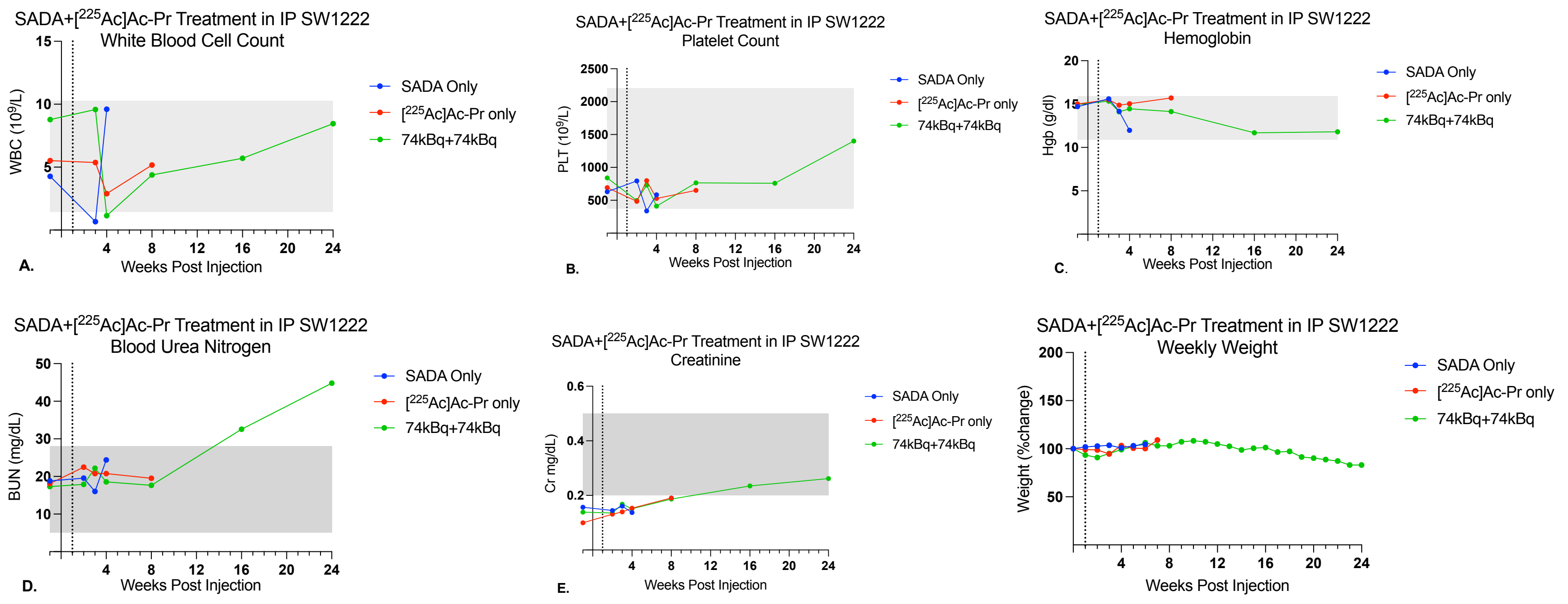


**Figure S10:** Representative hematology and renal markers from treatment of IP SW1222 model with SADA + [$^{225}$Ac]Ac-Pr. Groups were [$^{225}$Ac]Ac-Pr only, SADA only, GPA33-SADA treatment receiving 74kBq+74kBq. Mice were followed for a period of 24 weeks. **(A)** White blood cell count (WBC) demonstrating transient leukopenia in the treated group at week 4 post injection. **(B)** Platelet (Plt) demonstrating no period of thrombocytopenia. **(C)** Hemoglobin (Hgb) demonstrating no period of anemia. **(D)** Blood Urea Nitrogen (BUN), a maker of renal function, demonstrating an elevation in treatment group beginning at 16 weeks, correlating with histopathological evaluation. **(E)** Creatinine (Cr) demonstrating no change from baseline range throughout the study period. **(F)** Weights, measured as a % change from baseline at week 0, exhibiting stable weight over time in treated mice.

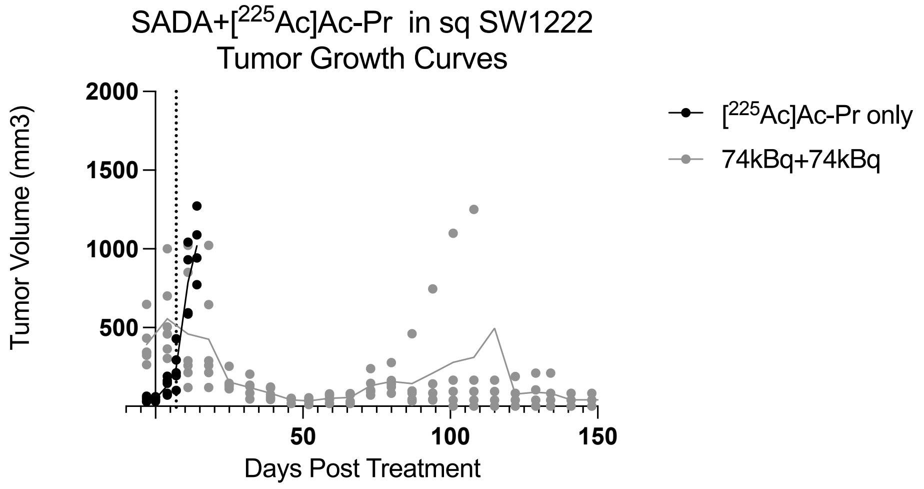


**Figure S11:** Tumor growth curves after treatment of subcutaneous SW1222 in an expanded cohort (n=10) in order to confirm treatment dose with GPA33-SADA + [$^{225}$Ac]Ac-Pr at a cycle of two doses 74kBq+74kBq 7 days apart. Mice were followed for 140 days, in which 4 mice were seen to have no tumor burden at the study endpoint. 6 mice had tumor recurrence observed at 90-100 days which followed a more indolent growth pattern than what was seen in untreated controls. Of note, starting tumor sizes were on average 200-500mm$^3$ larger than the standard experimental starting point.

| Treatment | Months post treatment (mouse age per group) | N | Renal pathology severity (1-4)* | Microscopic findings (# affected kidneys per animal / total) |
|---|---|---|---|---|
| SADA Only | 5 mon, 6 mon | 2 | 1-2 | Tubular attenuation, degeneration and/or regeneration (2/2) ; chronic interstitial nephritis (2/2) |
| 37kBq, single cycle | 3 mon (1), 5 mon (2), 6 mon (1) | 4 | 0-3& | Tubular attenuation (1/4) ; Hyaline Glomerulopathy (1/4) |
| 74kBq, single cycle | 3 mon (1), 5 mon (1), 6 mon (3) | 5 | 0 | Normal (0/5) |
| 37kBq+37KBq | 3 mon (1), 5 mon (1), 6 mon (3) | 5 | 1-3& | Tubular attenuation, degeneration, and basophilia, tubular single cell necrosis, tubular casts/proteinosis, and/or karyomegaly (4/5) ; Hyaline Glomerulopathy (2/5) |
| 37kBq+74KBq | 3 mon (1), 5 mon (1), 6 mon (2) | 4 | 1-2 | Tubular atrophy, degeneration and/or basophilia (2/4) ; chronic interstitial nephritis (2/4) |
| 74kBq+74KBq | 3 mon (1), 5 mon (2), 6 mon (2) | 5 | 1-4& | Tubular degeneration, attenuation, and/or tubular epithelial single cell necrosis (5/5)<br>Hyaline Glomerulopathy (3/5)<br>Lymphoma (1/5) |
| [$^{225}$Ac]Ac-Pr Only | 3 mon (1), 5 mon (1), 6 mon (3) | 5 | 1-3 | Chronic interstitial nephritis and/or fibrosis (3/5) |

* Renal pathology severity were minimal (1), mild (2), moderate (3) and severe (4)

**Table S1.** Renal histopathology in nontumor bearing athymic nude mice treated with dose escalating anti-GPA33 SADA SADA [$^{225}$Ac]Ac-Pr alpha-pretargeted radioimmunotherapy.

| Treatment | Months post treatment (mouse age per group) | N | Ovaries pathology severity (1-4)* | Microscopic findings (# affected organs per animal / total) |
|---|---|---|---|---|
| SADA Only | 5 mon, 6 mon | 5 | 0 | Normal |
| 37kBq, single cycle | 3 mon (1), 5 mon (2), 6 mon (1) | 4 | 0 | Normal |
| 74kBq, single cycle | 3 mon (1), 5 mon (1), 6 mon (3) | 5 | 0 | Normal |
| 37kBq+37KBq | 3 mon (1), 5 mon (1), 6 mon (3) | 5 | 2 | Cortical atrophy (3/5) |
| 37kBq+74KBq | 3 mon (1), 5 mon (1), 6 mon (2) | 4 | 2-4 | Cortical atrophy (2/4) |
| 74kBq+74KBq | 3 mon (1), 5 mon (2), 6 mon (2) | 5 | 1-4 | Cortical atrophy (3/5). Lymphoma (1/5) |
| [$^{225}$Ac]Ac-Pr Only | 3 mon (1), 5 mon (1), 6 mon (3) | 5 | 2-3 | Cortical atrophy (1/5) |

* The ovaries of one mouse was not available (NA).

**Table S2.** Ovarian histopathology in nontumor bearing athymic nude mice treated with dose escalating anti-GPA33 SADA SADA [$^{225}$Ac]Ac-Pr alpha-pretargeted radioimmunotherapy.

| Treatment | Months post treatment (mouse age per group) | N | Renal pathology severity (1-4)* | Microscopic findings (# affected kidneys per animal / total) |
|---|---|---|---|---|
| [$^{225}$Ac]Ac-Pr Only | 2 mon | 5 | 0-1 | Normal (4/5)<br>Focal tubular degeneration, Interstitial fibrosis (1/5) |
| SADA Only | 2 mon | 5 | 0-2 | Normal (2/5)<br>Tubular loss, degeneration and atrophy and interstitial fibrosis (1/5)<br>Tubular basophilia (1/5)<br>Tubular regeneration (1/5) |
| 37kBq | 3 mon | 5 | 0-1 | Normal (4/5)<br>Chronic interstitial nephritis (1/5) |
| 74kBq | 4 mon (3), 9 mon (1) | 4 | 0-3 | Normal (1/4)<br>Tubular degeneration, tubular epithelial single cell necrosis and/or karyomegaly (3/4)<br>Subintimal arterial mineralization (1/4) |
| 37kBq+37kBq | 4 mon (1), 9 mon (2), 13 mon (2) | 5 | 0-2 | Normal (1/5)<br>Tubular attenuation, degeneration, loss, mineralization, and/or casts/proteinosis (4/5) ;<br>Hyaline glomerulopathy (2/5); Chronic interstitial nephritis (2/5) |
| 37kBq+37kBq | 4 mon (1), 9 mon (3), 13 mon (1) | 5 | 0-3 | Tubular attenuations, degeneration, mineralization, regeneration, and/or casts/proteinosis (5/5)<br>Hyaline glomerulopathy (1/5); Chronic interstitial nephritis (1/5) |
| 74kBq+74kBq | 7mon (1), 13 mon (3) | 5 | 1-4 | Tubular attenuation, degeneration, loss, and casts/proteinosis (3/5); Interstitial fibrosis (3/5); Hyaline glomerulopathy (3/5)<br>Lymphoma (1/5) |
| 74kBq+74KBq, GD2 SADA | 2 mon | 5 | 0 | Normal (5/5) |

* Renal pathology severity were minimal (1), mild (2), moderate (3) and severe (4).
& Hyaline glomerulopathy has the highest pathology scores.

**Table S3.** Renal histopathology in athymic nude mice bearing sq SW1222 tumors treated with dose escalating anti-GPA33 SADA [$^{225}$Ac]Ac-Pr alpha-pretargeted radioimmunotherapy.

| Treatment | Months post treatment (mouse age per group) | N | Ovaries pathology severity (1-4)* | Microscopic findings (# affected organs per animal / total) |
|---|---|---|---|---|
| [$^{225}$Ac]Ac-Pr Only | 2 mon | 5 | 0 | Normal (5/5) |
| SADA Only | 2 mon | 5 | 0 | Normal (4/4*) |
| 37kBq | 3 mon | 4 | 0 | Normal (4/4) |
| 74kBq | 4 mon (3), 9 mon (1) | 4 | NA | NA |
| 37kBq+37kBq | 4 mon (1), 9 mon (2), 13 mon (2) | 4 | 2-3 | Cortical atrophy and/or follicular degeneration (2/2). Ovaries from 2 mice not collected |
| 37kBq+37kBq | 4 mon (1), 9 mon (3), 13 mon (1) | 5 | 3 | Cortical atrophy and/or follicular degeneration (1/1). Ovaries from 4 mice not collected |
| 74kBq+74kBq | 13 mon (3) | 5 | 2-4 | Cortical atrophy and/or degeneration (3/4).<br>Lymphoma (1/4)<br>Ovaries from 1 mouse not collected |
| 74kBq+74KBq, GD2 SADA | 2 mon | 5 | 0 | Normal (5/5) |

* The ovaries of one mouse was not available (NA).
* Ovarian pathology severity were minimal (1), mild (2), moderate (3) and severe (4).

**Table S4.** Ovarian histopathology in athymic nude mice bearing sq SW1222 tumors treated with dose escalating anti-GPA33 SADA [$^{225}$Ac]Ac-Pr alpha-pretargeted radioimmunotherapy.

| Treatment | Months post treatment (mouse age per group) | N | Renal pathology severity (1-4)* | Microscopic findings (# affected kidneys per animal / total) |
|---|---|---|---|---|
| SADA Only | | 4 | 1-3& | Normal (3/4), tubular basophilia (1/4); Interstitial mixed lymphocytic and neutrophilic infiltrates (1/4) |
| [$^{225}$Ac]Ac-Pr Only | | 5 | 0 | Normal (0/5) |
| 74kBq+74KBq | | 5 | 2-4 | Tubular attenuation, degeneration, tubular casts/proteinosis (5/5), and karyomegaly (5/5); Interstitial fibrosis (4/5); Hyaline glomerulopathy (5/5) |

* Renal pathology severity were minimal (1), mild (2), moderate (3) and severe (4).
& Hyaline glomerulopathy has the highest pathology scores.

**Table S5.** Renal histopathology in athymic nude mice bearing sq LS174T tumors treated with anti-GPA33 SADA [$^{225}$Ac]Ac-Pr alpha-pretargeted radioimmunotherapy radioimmunotherapy.

| Treatment | Months post treatment (mouse age per group) | N | Ovaries pathology severity (1-4)* | Microscopic findings (# affected organs per animal / total) |
|---|---|---|---|---|
| SADA Only | | 4 | 0 | Normal (1/1)* |
| [$^{225}$Ac]Ac-Pr Only | | 5 | 0 | Normal (2/2); Cortical atrophy (1/3)* |
| 74kBq+74KBq | | 5 | 4 | Cortical atrophy (2/2)* |

* The ovaries of the remaining mice were  not available (NA).

**Table S6.** Ovarian histopathology in athymic nude mice bearing sq LS174T tumors treated with anti-GPA33 SADA [$^{225}$Ac]Ac-Pr alpha-pretargeted radioimmunotherapy.

| Treatment | Months post treatment (Number of mice) | N | Renal pathology severity (1-4)* | Microscopic findings (# affected kidneys per animal / total) |
|---|---|---|---|---|
| SADA Only | 1 (1), 1.4 (1), 1.8 (1), 2 mon (1) | 4 | 0-1 | Normal (3/4)<br>Karyomegaly (1/4) |
| [$^{225}$Ac]Ac-Pr Only | 1 (1), 2 (2), 1.5 mon (1) | 4 | 0-1 | Normal (2/5)<br>Tubular cast (1/4)<br>Carcinoma infiltration (1/4) |
| 74kBq+74KBq | 2.5 (1), 3.3 (2), 4 (2), 4.6 (1), 5.5 (1), 6 mon (2) | 9 | 1-4 | Tubular attenuation, degeneration regeneration, loss, and casts/proteinosis (8/9); Tubular casts (1/9); Interstitial fibrosis (7/9); Hyaline glomerulopathy (4/9) |

* Renal pathology severity were minimal (1), mild (2), moderate (3) and severe (4).

**Table S7.** Renal histopathology in athymic nude mice bearing sq LOVO tumors treated with anti-GPA33 SADA [$^{225}$Ac]Ac-Pr alpha-pretargeted radioimmunotherapy.

| Treatment | Months post treatment (mouse age per group) | N | Ovaries pathology severity (1-4)* | Microscopic findings (# affected organs per animal / total) |
|---|---|---|---|---|
| SADA Only | 1 (1), 1.4 (1), 1.8 (1), 2 mon (1) | 4 | 0 | Normal (4/4) |
| [$^{225}$Ac]Ac-Pr Only | 1 (1), 2 (2), 1.5 mon (1) | 4 | 0 | Normal (3/4*) |
| 74kBq+74KBq | 2.5 (1), 3.3 (2), 4 (2), 4.6 (1), 5.5 (1), 6 mon (2) | 9 | 4 | Cortical atrophy and/or degeneration (9/9). |

* The ovaries of one mouse was not available
* Ovarian pathology severity were minimal (1), mild (2), moderate (3) and severe (4).

**Table S8.** Ovarian histopathology in athymic nude mice bearing sq LOVO tumors treated with anti-GPA33 SADA [$^{225}$Ac]Ac-Pr alpha-pretargeted radioimmunotherapy.

| Treatment | Months post treatment (mouse age per group) | N | Renal pathology severity (1-4)* | Microscopic findings (# affected kidneys per animal / total) |
|---|---|---|---|---|
| SADA Only | 1.5 mon | 2 | 0 | Normal (2/2) |
| [$^{225}$Ac]Ac-Pr Only | 1.5 - 3.5 mon | 4 | 2-3 | Normal (3/4); Tubular attenuation, degeneration, and karyomegaly, chronic nephritis, and hydronephrosis (1/4) |
| 74kBq+74KBq | 3 - 5 mon | 10 | 1-4 | Tubular attenuation, degeneration, tubular casts/proteinosis (10/10), and karyomegaly (9/10); Chronic nephritis (2/10); Interstitial fibrosis (2/10); Hyaline glomerulopathy (2/10) |

* Renal pathology severity were minimal (1), mild (2), moderate (3) and severe (4).
& Hyaline glomerulopathy has the highest pathology scores.

**Table S9.** Renal histopathology in athymic nude mice bearing ip SW1222-luc tumors treated with anti-GPA33 SADA [$^{225}$Ac]Ac-Pr alpha-pretargeted radioimmunotherapy**.**

| Treatment | Months post treatment (mouse age per group) | N | Ovarian pathology severity (1-4)* | Microscopic findings (# affected organs per animal / total) |
|---|---|---|---|---|
| SADA Only | 1.5 mon | 2 | 0 | Normal (1/2)* |
| [$^{225}$Ac]Ac-Pr Only | 1.5 - 3.5 mon | 4 | 0 | Normal (4/4) |
| 74kBq+74KBq | 3 - 5 mon | 10 | 3-4 | Cortical atrophy with follicular loss and/or degeneration (7/10)* |

* The ovaries of the remaining mice were not available.

**Table S10.** Ovarian histopathology in athymic nude mice bearing ip SW1222-luc tumors treated with anti-GPA33 SADA [$^{225}$Ac]Ac-Pr alpha-pretargeted radioimmunotherapy.
